# Simeprevir potently suppresses SARS-CoV-2 replication and synergizes with remdesivir

**DOI:** 10.1101/2020.05.26.116020

**Authors:** Ho Sing Lo, Kenrie Pui Yan Hui, Hei-Ming Lai, Khadija Shahed Khan, Simranjeet Kaur, Junzhe Huang, Zhongqi Li, Anthony K. N. Chan, Hayley Hei-Yin Cheung, Ka-Chun Ng, John Chi Wang Ho, Yu Wai Chen, Bowen Ma, Peter Man-Hin Cheung, Donghyuk Shin, Kaidao Wang, Meng-Hsuan Lee, Barbara Selisko, Cecilia Eydoux, Jean-Claude Guillemot, Bruno Canard, Kuen-Phon Wu, Po-Huang Liang, Ivan Dikic, Zhong Zuo, Francis K. L. Chan, David S. C. Hui, Vincent C. T. Mok, Kam-Bo Wong, Ho Ko, Wei Shen Aik, Michael Chi Wai Chan, Wai-Lung Ng

## Abstract

The outbreak of coronavirus disease 2019 (COVID-19), caused by the severe acute respiratory syndrome coronavirus 2 (SARS-CoV-2), is a global threat to human health. Using a multidisciplinary approach, we identified and validated the hepatitis C virus (HCV) protease inhibitor simeprevir as an especially promising repurposable drug for treating COVID-19. Simeprevir potently reduces SARS-CoV-2 viral load by multiple orders of magnitude and synergizes with remdesivir *in vitro*. Mechanistically, we showed that simeprevir inhibits the main protease (M^pro^) and unexpectedly the RNA-dependent RNA polymerase (RdRp). Our results thus reveal the viral protein targets of simeprevir, and provide preclinical rationale for the combination of simeprevir and remdesivir for the pharmacological management of COVID-19 patients.

**One Sentence Summary:** Discovery of simeprevir as a potent suppressor of SARS-CoV-2 viral replication that synergizes with remdesivir.

## Introduction

The recent outbreak of infection by the novel betacoronavirus severe acute respiratory syndrome coronavirus 2 (SARS-CoV-2) has spread to almost all countries and claimed more than 760,000 lives worldwide (WHO situation report 209, August 16, 2020). Alarming features of COVID-19 include a high risk of clustered outbreak both in community and nosocomial settings, and up to one-fifth severe/critically ill proportion of symptomatic inpatients reported^1–4^. Furthermore, a significant proportion of infected individuals are asymptomatic, substantially delaying their diagnoses, hence facilitating the widespread dissemination of COVID-19^5^. With a dire need for effective therapeutics that can reduce both clinical severity and viral shedding, numerous antiviral candidates have been under clinical trials or in compassionate use for the treatment of SARS-CoV-2 infection^6^.

Several antivirals under study are hypothesized or proven to target the key mediator of a specific step in the SARS-CoV-2 viral replication cycle. For instance, lopinavir/ritonavir (LPV/r) and danoprevir have been proposed to inhibit the SARS-CoV-2 main protease (M^pro^, also called 3CL^Pro^) needed for the maturation of multiple viral proteins; chloroquine (CQ) / hydroxychloroquine (HCQ) [alone or combined with azithromycin (AZ)] may abrogate viral replication by inhibiting endosomal acidification crucial for viral entry^7,8^; nucleoside analogues such as remdesivir, ribavirin, favipiravir and EIDD-2801 likely inhibit the SARS-CoV-2 nsp12 RNA-dependent RNA polymerase (RdRp) and/or induce lethal mutations during viral RNA replication^9–11^. Unfortunately, on the clinical aspect, LPV/r failed to demonstrate clinical benefits in well-powered randomized controlled trials (RCTs), while HCQ and/or AZ also failed to demonstrate benefits in observational studies^12–14^. Meanwhile, LPV/r, CQ/HCQ and AZ may even increase the incidence of adverse events^14–16^. Although remdesivir is widely considered as one of the most promising candidates, latest RCTs only revealed marginal shortening of disease duration in patients treated^17^. Therefore, further efforts are required to search for more potent, readily repurposable therapeutic agents for SARS-CoV-2 infection, either as sole therapy or in combination with other drugs to enhance their efficacy.

Ideally, the candidate drugs need to be readily available as intravenous and/or oral formulation(s), possess favourable pharmacokinetics properties as anti-infectives, and do not cause adverse events during the treatment of SARS-CoV-2 infection (e.g. non-specific immunosuppression, arrhythmia or respiratory side effects). Two complementary approaches have been adopted to identify novel drugs or compounds that can suppress SARS-CoV-2 replication. One approach relies on *in vitro* profiling of the antiviral efficacy of up to thousands of compounds in early clinical development, or drugs already approved by the U.S. Food and Drug Administration (FDA)^18–22^. On the other hand, as the crystal structure of the M^pro^ ^23,24^, papain-like protease (PL^pro^)^25^ and the cryo-EM structure of the nsp12-nsp7-nsp8 RdRp complex^11,26^ of the SARS-CoV-2 virus became available, the structure-based development of their specific inhibitors becomes feasible. Structure-aided screening will enable the discovery of novel compounds as highly potent inhibitors^27^ as well as the repurposing of readily available drugs as anti-CoV agents for fast-track clinical trials.

Here we report our results regarding the discovery of FDA-approved drugs potentially active against the SARS-CoV-2. *In vitro* experiments led to the identification of simeprevir, a hepatitis C virus (HCV) NS3A/4 protease inhibitor^28^, as a potent inhibitor of SARS-CoV-2 replication (**Fig. 1A**). Importantly, simeprevir acts synergistically with remdesivir, whereby the effective dose of remdesivir could be lowered by multiple-fold by simeprevir at physiologically feasible concentrations. Interestingly, biochemical and molecular characterizations revealed that simeprevir inhibits both the M^pro^ protease and RdRp polymerase activities. This unexpected anti-SARS-CoV-2 mechanism of simeprevir provides hints on novel antiviral strategies.

**Fig. 1.**
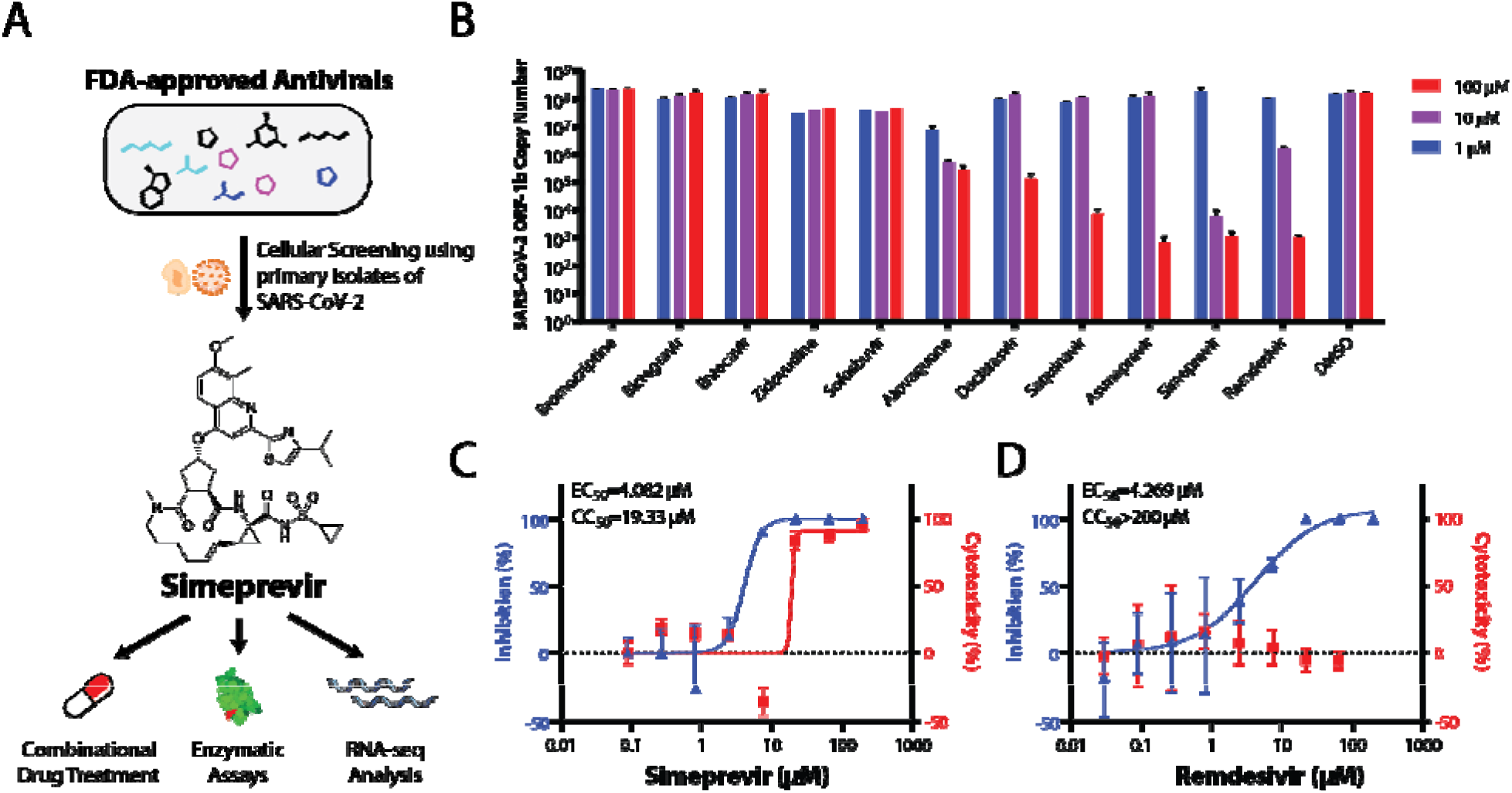
Repurposing FDA-approved Drugs for SARS-CoV-2 through cellular screening. **(A)** Summar of methodology used in this paper. **(B)** Screening for FDA-approved small molecule therapeutics for activities in suppressing SARS-CoV-2 replication in Vero E6 cells. Dose-response curves in the suppression of SARS-CoV-2 replication in Vero E6 cells and cytotoxicity for simeprevir **(C)** and remdesivir **(D)** are shown. Data points in all plots represent mean ± S.E.M‥ For all data points, *n* = 3 replicates.

## Results

### A prioritized screening identifies simeprevir as a potent suppressor of SARS-CoV-2 replication in a cellular infection model

Given our goal of identifying immediately usable and practical drugs against SARS-CoV-2, we prioritized a list of repurposing drug candidates for *in vitro* testing based on joint considerations on safety, pharmacokinetics, drug formulation availability, and feasibility of rapidly conducting clinical trials (**Table S1**). We focused on FDA-approved antivirals (including simeprevir, saquinavir, daclatasvir, ribavirin, sofosbuvir and zidovudine), and drugs whose primary indication was not antiviral but had reported antiviral activity (including bromocriptine and atovaquone). Remdesivir was also tested for comparison of efficacy and as a positive control.

In a Vero E6 cellular infection model, we found the macrocyclic HCV protease inhibitor simeprevir as the only prioritized drug candidate that showed potent suppression of SARS-CoV-2 replication in the ≤ 10 μM range (**Fig. 1B**). More detailed dose-response characterization found that simeprevir has a potency comparable to remdesivir (**Fig. 1C, D**). The half-maximal effective concentration (EC50) of simeprevir was determined to be 4.08 μM, while the 50% cytotoxicity concentration (CC50) was 19.33 μM (**Fig. 1C, D**). In a physiologically relevant human lung epithelial cell model, ACE2-expressing A549 cells (A549-ACE2) infected with SARS-CoV-2, we also observed the strong antiviral effect of simeprevir^29^ (**Supplementary Fig. 1**). The cytotoxicity data are also in line with the reported *in vitro* and *in vivo* safety pharmacological profiling using human cell lines, genotoxicity assays, and animal models^28,30^. These data suggest that a desirable therapeutic window exists for the suppression of SARS-CoV-2 replication with simeprevir.

### Simeprevir potentiates the suppression of SARS-CoV-2 replication by remdesivir

While simeprevir is a potential candidate for clinical use alone, we hypothesized that it may also have a synergistic effect with remdesivir, thereby mitigating its reported adverse effects, improving its efficacy, and broadening its applicability^17^. Indeed, combining simeprevir and remdesivir at various concentrations apparently provided much greater suppression of SARS-CoV-2 replication than remdesivir alone, while they did not synergize to increase cytotoxicity (**Fig. 2A**). Importantly, such effects were not merely additive, as the excess over Bliss score suggested synergism at 3.3 μM simeprevir and 1.1 – 10 μM remdesivir in suppressing SARS-CoV-2 replication (**Fig. 2B**).

**Fig. 2.**
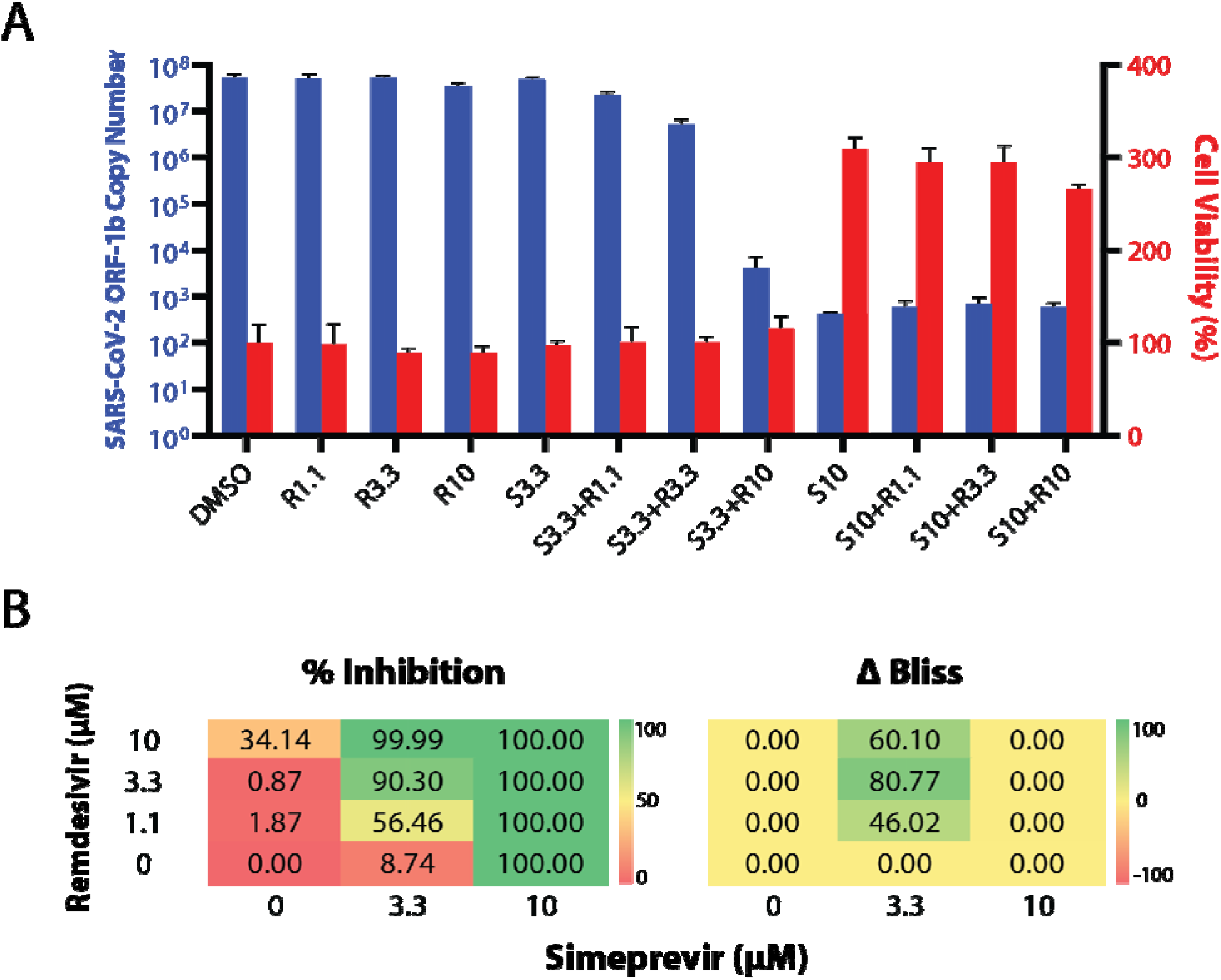
**(A)** Viral replication-suppression efficacies of different combinations of simeprevir and remdesivir concentrations. The numbers after S (simeprevir) and R (remdesivir) indicate the respective drug concentrations in μM. Data points in all plots represent mean ± S.E.M‥ For all data points, *n* = 3 replicates. **(B)** Bliss score analyses of synergism. Left panel: Diagram showing 12 combinations of simeprevir and remdesivir and their respective percentage inhibition (% inhibition, color-coded) of SARS-CoV-2 replication in Vero E6 cells compared to DMSO controls. Right panel: Excess over Bliss score (ΔBliss, color-coded) of different drug combinations. A positive and negative number indicates a likely synergistic and antagonistic effect, respectively, while a zero value indicates independence of action.

### Simeprevir weakly inhibits the M^pro^ and RdRp but does not inhibit PL^pro^ at physiologically feasible concentrations

The desirable anti-SARS-CoV-2 effect of simeprevir prompted us to determine its mechanism of action. Given that simeprevir is an HCV NS3/4A protease inhibitor, we first investigated its inhibitory activity against SARS-CoV-2 M^pro^ and PL^pro^ using biochemical assays^31,32^ (**Fig. 3**). We found inhibition of M^pro^ by simeprevir with half-maximal inhibitory concentration (IC_50_) of 9.6 ± 2.3 μM (**Fig. 3A**), two times higher than the EC_50_ determined from our cell-based assay. The substrate cleavage was further verified with SDS-PAGE (**Supplementary Fig. 3**). Docking simeprevir against the apo protein crystal structure of SARS-CoV-2 M^pro^ suggested a putative binding mode with a score of −9.9 kcal mol^−1^ (**Supplementary Fig. 4**). This binding mode is consistent with a recent docking study using a homology model of SARS-CoV-2 M^pro^ ^33^. On the other hand, no inhibition of PL^pro^ activity was observed at physiologically feasible concentrations of simeprevir, with either ISG15 or ubiquitin as substrate (**Fig. 3B, Supplementary Fig. 5, 6**).

**Fig. 3.**
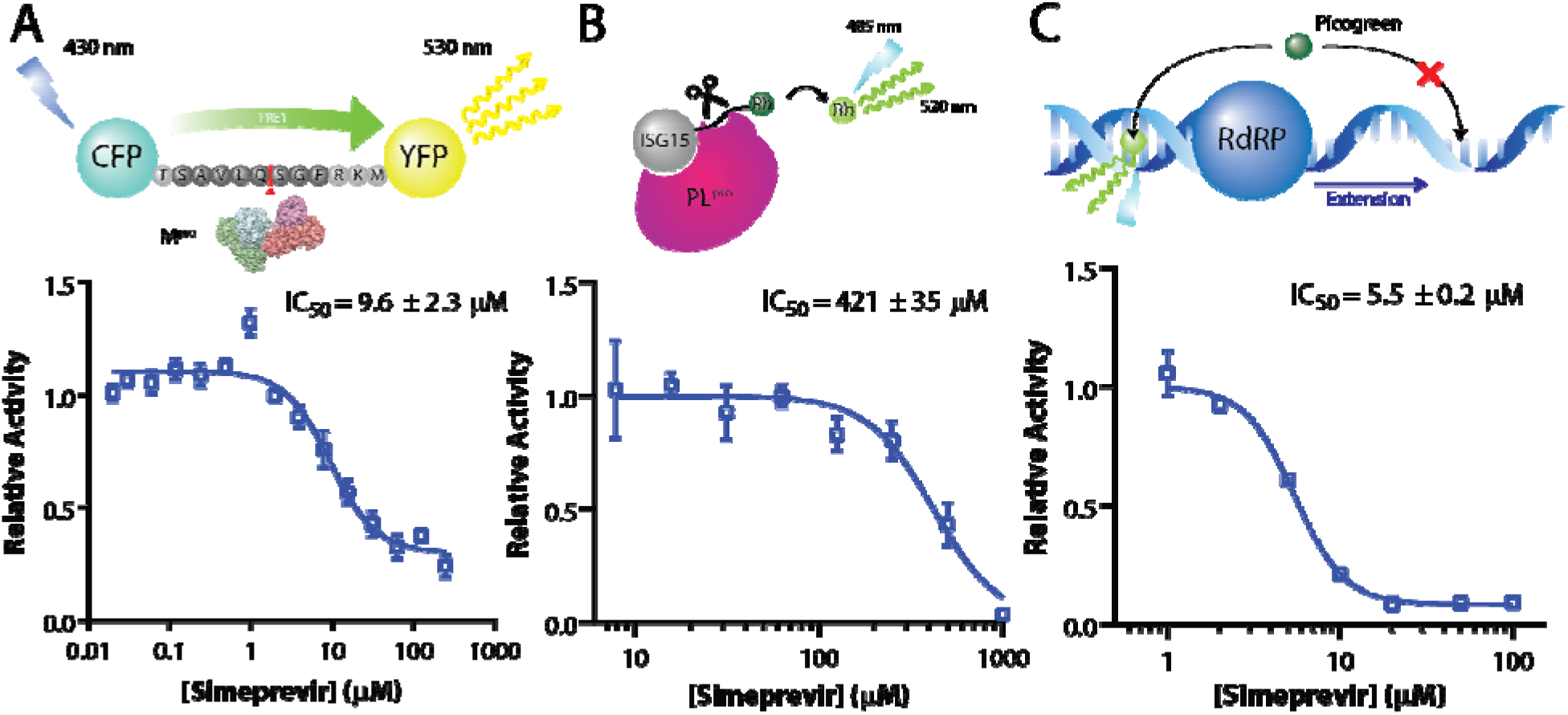
Simeprevir weakly inhibits M^pro^ and RdRp. Assay scheme and enzyme activity of main protease (M^pro^), papain-like protease (PL^pro^) and RNA-dependent RNA polymerase (RdRp) under variou concentrations of simeprevir. **(A)** For M^pro^, a CFP-YFP conjugate with a M^pro^ cleavage site linker is utilized, where relative activity is determined by the residual FRET efficiency after cleavage. **(B)** For PL^pro^, rhodamine-conjugated ISG15 is used as a substrate for the enzyme, whose relative activity is determined by release of the fluorophore. **(C)** For RdRp, an extension assay based on the dsRNA-binding property of the intercalating agent Picogreen was established. Data points in all plots represent mean ± S.E.M‥ For all data points, *n* = 3 replicates.

We speculated that the weak inhibition of M^pro^ protease activity by simeprevir could not fully account for its antiviral effect towards SARS-CoV-2. To identify additional target(s), we next docked simeprevir alongside several nucleoside analogues (remdesivir, ribavirin, and favipiravir) against the motif F active site of the cryo-EM structure of the SARS-CoV-2 nsp12 RdRp (**Supplementary Fig. 7A**). Interestingly, the docking results revealed that simeprevir had a higher binding score than the nucleoside analogues (**Supplementary Fig. 7B**). To test this experimentally, we established and performed RdRp primer extension assays using recombinant nsp12, nsp7, and nsp8 of SARS-CoV^34^. Intriguingly, simeprevir showed low micromolar-range inhibition towards SARS-CoV RdRp as validated by both a gel-based assay (**Supplementary Fig. 8**) and a Picogreen fluorescence-based assay, with an IC_50_ value of 5.5 ± 0.2 μM (**Fig. 3C**). Collectively, the assay data suggested that simeprevir inhibits the enzymatic activities of both M^pro^ and RdRp but not PL^pro^.

### RNA Sequencing identifies significant downregulation of viral defense responses upon the treatment of simeprevir

While inhibition of viral targets seems to be a primary mechanism of action of simeprevir, the weak inhibitory effects observed in biochemical assays as well as its cytotoxicity suggest the possibility of additional host-mediated antiviral response. To further elucidate the antiviral mechanism of simeprevir, we next performed RNA sequencing on Vero E6 cells to reveal the transcriptomic changes upon drug treatment (**Fig. 4A**). In line with the literature, SARS-CoV-2 infection induced type I interferon and chemokine response (**Supplementary Fig. 9**)^35,36^. In mock-infected cells, simeprevir treatment (at 1.1 μM or 3.3 μM) did not induce any significant changes of differentially expressed genes (DEGs); while in SARS-CoV-2-infected cells, a small number of DEGs was observed (**Fig. 4B**). Gene set enrichment analyses (GSEA) of infected cells using Reactome gene sets revealed significant positive enrichment of 93 gene sets in the simeprevir-treated samples, including histone lysine/arginine methylation, histone demethylation, and cell cycle control (**Fig. 4C, Supplementary table 2**). On the other hand, gene sets with the gene ontology (GO) terms “defense response to virus” and “response to type I interferon” were negatively enriched, suggesting overall downregulation of these gene sets (**Fig. 4D**). In the latter set, green monkey orthologs of crucial human innate immune-related genes (e.g. IFIT1-3, USP18) and interferon-stimulated genes (e.g. ISG15) were downregulated with simeprevir treatment in a dose-dependent manner (**Fig. 4E**). Similarly, the downregulation of some of these genes as well as the proinflammatory cytokines IL-6 and interferon IFNL1 were also observed with remdesivir treatment (**Fig. 4F**).

**Fig. 4.**
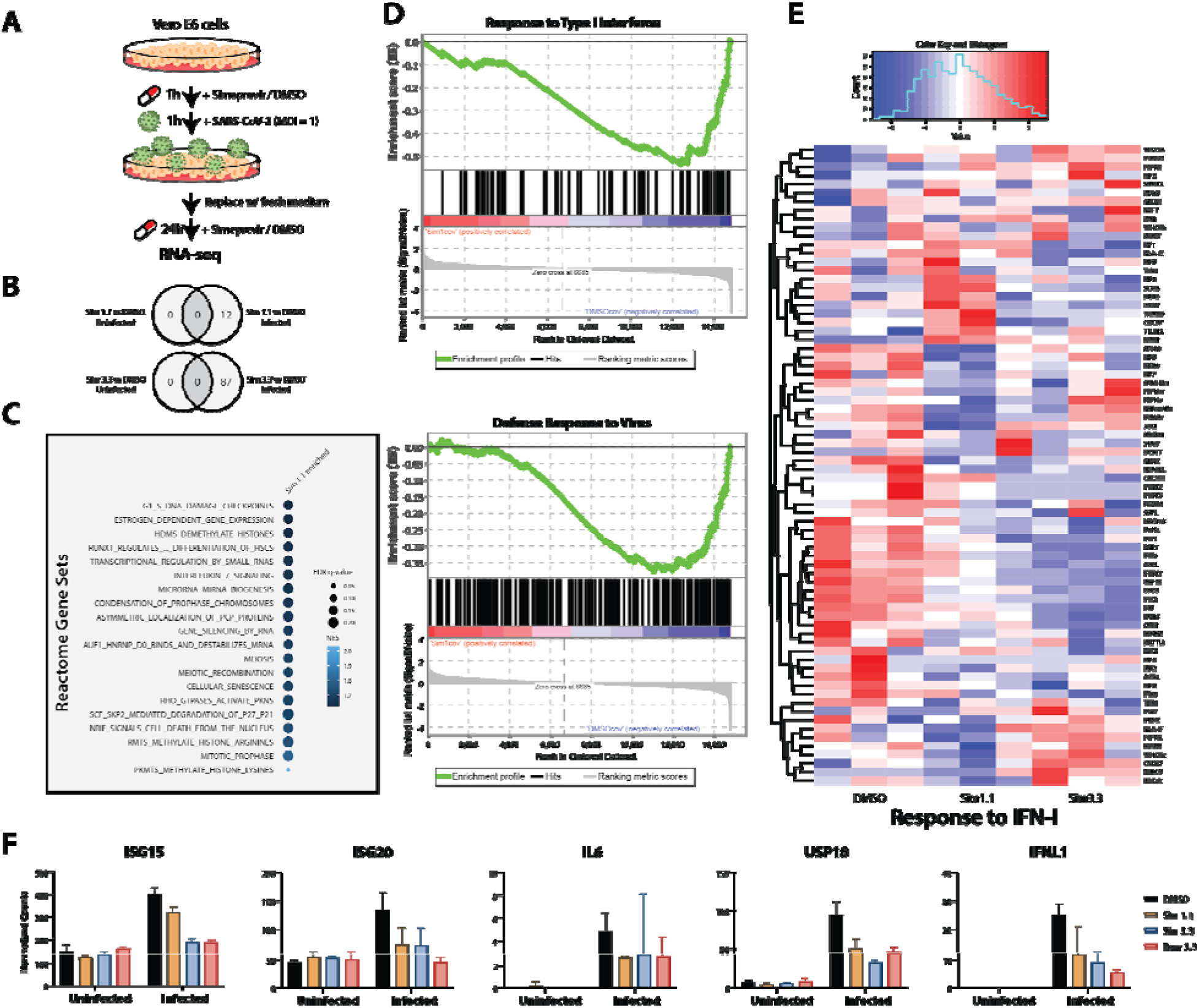
RNA-seq analysis of simeprevir-mediated host response and antiviral activity. **(A)** Schematic representation of RNA-seq sample preparation. Treatment sequence and incubation time of simeprevir and SARS-CoV-2 was indicated with arrows and legends. **(B)** Venn diagrams showing differentially expressed genes (DEGs) comparing simeprevir-treated (1.1μM or 3.3μM), infected and mock-infected samples. **(C)** Bubble plot of top 20 hits of positively enriched reactome gene sets under simeprevir treatment using gene set analysis (GSEA). Enriched gene sets were filtered with criteria false discovery rate (FDR) q-value < 0.25 and nominal p-value < 0.05 before ranked with their normalized enrichment scores (NES). **(D)** Enrichment plots of GSEA results using gene ontology (GO) gene sets. **(E)** Clustered heatmap showing the row-normalized expression level of genes belonging to GO term “response to type I interferon”. **(F)** Bar charts showing normalized counts from RNA-seq data of five genes involved in antiviral response. Data points in this plot represent mean ± S.E.M. For all data points, *n* = 3 replicates.

## Discussion

The novel coronavirus SARS-CoV-2 has gone from an emerging infection to a global pandemic with its high transmissibility. As human activities are becoming more aggressive and damaging to nature, future coronavirus pandemics are bound to happen. It is therefore essential to reduce the casualties by effective pharmacological management. Our study has successfully identified the readily repurposable, clinically practical antiviral simeprevir that could target SARS-CoV-2. Specifically, we found up to fourth-order suppression of viral genome copies by simeprevir at ≤ 10 μM in cell-based viral replication assays - a concentration that is expected to be attainable in human lung tissues with ≥ 150mg daily dosing based on available pharmacokinetic data^37,38^.

In addition, we discovered that simeprevir can synergize with remdesivir in inhibiting SARS-CoV-2 replication in a cellular model, potentially allowing lower doses of both drugs to be used to treat COVID-19. In a global pandemic with patients having diverse clinical characteristics, providing additional therapeutic options to remdesivir will be important to treat those who are intolerant or not responding to the drug ^17^, which can easily amount to tens of thousands of patients. As there is only one confirmed and approved therapy for COVID-19, a potentially repurposable drug can be rapidly tested in animal models before clinical trials to prepare for supply shortages or when remdesivir-resistant mutations arise. Combination treatment, such as simeprevir-remdesivir, may also help to reduce the dose required to alleviate side effects.

We note, however, there are also several limitations of simeprevir and the proposed simeprevir-remdesivir combination. Simeprevir requires dose adjustments in patients with Child-Pugh Class B or C cirrhosis, as well as in patients with East Asian ancestry^37^. In addition, simeprevir has been taken off the market since 2018 due to the emergence of next-generation HCV protease inhibitors, hence its supply may not be ramped up easily. Noteworthily, simeprevir is metabolized by the CYP3A4 enzyme with saturable kinetics^37^ while remdesivir itself is not only a substrate of CYP3A4 but also a CYP3A4 inhibitor. Whether such theoretical pharmacokinetic interaction will exacerbate liver toxicity or provide additional pharmacokinetic synergy (in addition to pharmacodynamic synergy) *in vivo* remains to be tested.

Mechanistically, we found that simeprevir suppresses SARS-CoV-2 replication by targeting at least two viral proteins – it weakly inhibits M^pro^ at ~10 μM and unexpectedly inhibits RdRp at ~5 μM. The potency towards M^pro^ is consistent with the IC_50_ of ~13.7 μM as determined in a parallel study^39^. Our gel-based assay (**Supplementary Fig. 9**) suggested that simeprevir interferes with RNA-binding of RdRp because less probe was extended but to full length. This is also supported by the *in silico* docking results, in which simeprevir is docked to a highly conserved RNA binding site showing no amino acid polymorphism between SARS-CoV and SARS-CoV-2 (**Supplementary Fig. 8A**). This putative binding mode hints that simeprevir might block the RNA binding site while remdesivir might target the nucleoside entry site, potentially resulting in a synergistic effect. Importantly, the high similarity in sequence (96% identity) and structure between SARS-CoV-2 and SARS-CoV also suggest that simeprevir might act as a broad-spectrum antiviral in the *Coronaviridae* family.

Furthermore, the discrepancy between RdRp and M^pro^ inhibitory potency versus *in vitro* inhibitory potency of SARS-CoV-2 replication suggested additional mechanism(s) of action of simeprevir. Our RNA-seq and GSEA analysis revealed several molecular pathways that warrant future investigation. A possible direction is immune modulation *via* epigenetic regulations, which could mediate viral infection (*via* SWI/SNF chromatin remodeling complex and histone H3.3 complex)^40^, interferon-induced antiviral response (*via* H3K79 methylation)^41^, and host immune evasion (*via* alteration of DNA methylome)^42^. Whether the downregulation of type I IFN-related genes stems directly from simeprevir’s action or a reduction in viral load remains an open question. Collectively, simeprevir targets two viral proteins but may also act on the host proteins to suppress SARS-CoV-2 replication.

Given that simeprevir is originally a non-nucleoside antiviral targeting HCV protease, its inhibition towards RdRp is largely unexpected and represents a novel mechanism of action. Simeprevir thus holds promise to be a lead compound for the future development of dual inhibitors of M^pro^ and RdRp. It should be noted that the potencies of M^pro^ and RdRp inhibition may not entirely account for the strong suppression of SARS-CoV-2 viral replication. Therefore, further investigation of the mechanism of action of simeprevir may uncover new druggable targets for inhibiting SARS-CoV-2 replication.

## Materials and Methods

### Chemicals and reagents

Bromocriptine mesylate (BD118791), saquinavir (BD150839), bictegravir (BD767657), atovaquone (BD114807) and asunaprevir (BD626409) were purchased from BLD Pharmatech (Shanghai, China). Entecavir (HY-13623), zidovudine (HY-17413), sofosbuvir (HY-15005), daclatasvir (HY-10466), simeprevir (HY-10241), remdesivir (HY-104077) and remdesivir triphosphate sodium (HY-126303A) were purchased from MedChemExpress (Monmouth Junction, NJ). Drug stocks were made with DMSO.

### In vitro SARS-CoV-2 antiviral tests

SARS-CoV-2 virus (BetaCoV/Hong Kong/VM20001061/2020, SCoV2) was isolated from the nasopharyngeal aspirate and throat swab of a COVID-19 patient in Hong Kong using Vero E6 cells (ATCC CRL-1586). Vero E6 or A549-ACE2 cells were infected with SCoV2 at a multiplicity of infection (MOI) of 0.05 or 0.5, respectively, in the presence of varying concentrations and/or combinations of the test drugs. DMSO as the vehicle was used as a negative control. Antiviral activities were evaluated by quantification of SARS-CoV-2 ORF1b copy number in the culture supernatant by using quantitative real-time RT-PCR (qPCR) at 48 h post-infection with specific primers targeting the SARS-CoV-2 ORF1b^43^.

### In vitro drug cytotoxicity assays

*In vitro* cytotoxicity of the tested drugs was evaluated using thiazolyl blue tetrazolium bromide (MTT, Sigma-Aldrich)-based cell viability assays. Vero E6 cells were seeded onto 48-well plates and treated with indicated concentrations of simeprevir and/or remdesivir for 48 h. Treated cells were incubated with DMEM supplemented with 0.15 mg/mL MTT for 2 h, and formazan crystal products were dissolved with DMSO. Cell viability was quantified with colorimetric absorbance at 590 nm.

### Molecular docking simulations

Three-dimensional representations of chemical structures were extracted from the ZINC15 database (http://zinc15.docking.org)^44^, with the application of three selection filters –– Protomers, Anodyne, and Ref. ZINC15 subset DrugBank FDA (http://zinc15.docking.org./catalogs/dbfda/) were downloaded as the mol2 file format. The molecular structures were then converted to the pdbqt format (the input file format for AutoDock Vina) using MGLTools2-1.1 RC1 (sub-program “prepare_ligand”) (http://adfr.scripps.edu/versions/1.1/downloads.html). AutoDock Vina v1.1.2 was employed to perform docking experiments^45^. Docking of simeprevir on SARS-CoV-2 M^pro^ was performed with the target structure based on an apo protein crystal structure (PDB ID: 6YB7); the A:B dimer was generated by crystallographic symmetry. Docking was run with the substrate-binding residues set to be flexible. Docking of simeprevir and other active triphosphate forms of nucleotide analogues was performed against the nsp12 portion of the SARS-CoV-2 nsp12-nsp7-nsp8 complex cryo-EM structure (PDB ID: 6M71).

### Expression and purification of M^pro^ and its substrate for FRET assay

The sequence of SARS-CoV-2 M^pro^ was obtained from GenBank (accession number: YP_009725301), codon-optimized, and ordered from GenScript. A C-terminal hexahistidine-maltose binding protein (His_6_-MBP) tag with two in-between Factor Xa digestion sites were inserted. Expression and purification of SARS-CoV-2 M^pro^ was then performed as described for SARS-CoV M^pro^ ^31^. The protein substrate, where the cleavage sequence “TSAVLQSGFRKM” of M^pro^ was inserted between a cyan fluorescent protein and a yellow fluorescent protein, was expressed and purified as described^31^.

### In vitro M^pro^ inhibition assay

The inhibition assay was based on fluorescence resonance energy transfer (FRET) using a fluorescent protein-based substrate previously developed for SARS-CoV M^pro^ ^31,46^. 0.1 μM of purified SARS-CoV-2 M^pro^ was pre-incubated with 0 - 250 μM simeprevir in 20 mM HEPES pH 6.5, 120 mM NaCl, 0.4 mM EDTA, 4 mM DTT for 30 min before the reaction was initiated by addition of 10 μM protein substrate^32^. Protease activity was followed at 25 °C by FRET with excitation and emission wavelengths of 430 nm and 530 nm, respectively, using a multi-plate reader as previously described^31,46^. Reduction of fluorescence at 530 nm was fitted to a single exponential decay to obtain the observed rate constant (*k_obs_*). Relative activity of M^pro^ was defined as the ratio of *k_obs_* with inhibitors to that without. The relative IC_50_ value of simeprevir was determined by fitting the relative activity at different inhibitor concentration to a four-parameter logistics equation.

### Expression and purification of SARS-CoV nsp7L8, nsp12

The fusion protein nsp7-nsp8 (nsp7L8) was generated by inserting a GSGSGS linker sequence between the nsp7 and nsp8 coding sequences^34^. The nsp7L8, nsp8 and nsp12 were produced and purified independently as described^47^. The complex, referred to as the replication/transcription complex (RTC), was reconstituted with a 1:3:3 ratio of nsp12:nsp7L8:nsp8.

### In vitro RdRp inhibition assay - fluorescence-based

The assay was performed as previously described^48^. The compound concentration leading to a 50% inhibition of RTC-mediated RNA synthesis was determined as previously described. Briefly, poly(A) template and the SARS-CoV RTC was incubated 5 min at room temperature and then added to increasing concentration of compound. Reaction was started by adding UTP and incubated 20 min at 30°C. Reaction assays were stopped by the addition of 20 μl EDTA 100 mM. Positive and negative controls consisted of a reaction mix with 5% DMSO (final concentration) or EDTA (100 mM) instead of compounds, respectively. Picogreen® fluorescent reagent was diluted to 1/800 final in TE buffer according to the data manufacturer and aliquots were distributed into each well of the plate. The plate was incubated for 5 min in the dark at room temperature and the fluorescence signal was then read at 480 nm (excitation) and 530 nm (emission) using a TecanSafire2 microplate reader. IC_50_ was determined using the following equation: % of active enzyme = 100/(1+(I)^2^/IC_50_), where I is the concentration of inhibitor and 100% of activity is the fluorescence intensity without inhibitor. IC_50_ was determined from curve-fitting using the GraphPad Prism 8.

### In vitro RdRp inhibition assay - gel-based

Enzyme mix (10 μM nsp12, 30 μM nsp7L8, 30 μM nsp8) in complex buffer (25 mM HEPES pH 7.5, 150 NaCl, 5 mM TCEP, 5 mM MgCl_2_) was incubated for 10 min on ice and then diluted with reaction buffer (20 mM HEPES pH 7.5, 50 mM NaCl, 5 mM MgCl_2_) to 2 μM nsp12 (6 μM nsp7L8 and nsp8) to a final volume of 10 μl. The resulting enzyme complex was mixed with the 10 μl of 0.8 μM primer/ template (P/T) carrying a Cy5 fluorescent label at the 5’ end (P:Cy5-GUC AUU CUC C, T: UAG CUU CUU AGG AGA AUG AC) in reaction buffer, and incubated at 30°C for 10 min. Inhibitor was added in 2 μl to the elongation complex and reactions were immediately started with 18 μl of NTP mix in the reaction buffer. Final concentrations in the reactions were 0.5 μM nsp12 (1.5 μM nsp7L8 and nsp8), 0.2 μM P/T, 50 μM NTPs and the given concentrations of inhibitors. Samples of 8 μl were taken at given time points and mixed with 40 μl of formamide containing 10 mM EDTA. Ten-μl samples were analyzed by denaturing PAGE (20 % acrylamide, 7 M urea, TBE buffer); and product profiles visualized by a fluorescence imager (Amersham Typhoon). Quantification of product bands and analysis were performed using ImageQuant and Excel.

### In vitro PL^pro^ inhibition assay - fluorescence-based

The purification and assay of PL^pro^ activity was adapted from as previously described^25^. Briefly, a ISG15-C-term protein tagged with rhodamine was used as a substrate for the enzymatic assay. Because of the solubility of Simeprevir, we used an optimized reaction buffer (5% DMSO, 15% PEG300, 50 mM Tris-HCl (pH 7.5), 50 mM NaCl, 5 mM DTT). 5 uL of solution containing 0 - 2 mM of Simeprevir and 2 μM of ISG15c-rhodamine were aliquoted into a 384-well plate. Reaction was initiated by addition of 5 μL of 40 nM PL^pro^ to the well. Initial velocities of rhodamine release (36 - 240 seconds) were normalized against DMSO control. Reactions were conducted for 12 minutes with monitoring of fluorescence intensity at 485/520 nm using a microplate reader (PHERAstar FSX, BMG Labtech). The same experiment was repeated with ubiquitin-rhodamine substrate.

### In vitro PL^pro^ inhibition assay - Gel-based

All proteins used for ubiquitination and PL^pro^ biochemical assays were expressed in *E. coli* RIL cells with 0.6 mM IPTG over-induction at 16 °C. Rsp5 WW3-HECT with 2 mutations (Q808M, E809L) at the C-terminus was expressed using GST-fusion affinity tag followed by TEV protease digestion and purification by size exclusion chromatography. His-tagged PL^pro^, UBA1, UBCH7, ubiquitin K63R with extra cysteine at the N-terminus were also expressed and purified similarly, except hexahistidine tag used. Ubiquitin was further cross-linked with fluorescein-5-maleimide (Anaspec, Fremont, CA, US) and the poly-ubiquitination sample was generated following previous protocol ^49^. Deubiquitination assays using PL^pro^ (SARS-CoV-2) were carried out at 37 °C for 10 minutes using 1-100 μM Simeprevir and 1 μM PL^pro^. Final DMSO concentration of each reaction is 2%. The reaction was quenched by SDS sample buffer and analyzed by 4-20% SDS-gel (GenScript, Piscataway, NJ, US). Fluorescent ubiquitin signals were imaged using Thermo iBright exposed for 750 ms.

### Sample preparation for RNA-seq

Approximately 4 × 10^5^ Vero E6 cells were seeded onto each well of 12-well plates, in DMEM supplemented with 2% FBS, 4.5 g/L D-glucose, 4 mM L glutamine, 25 mM HEPES and 1% penicillin/streptomycin. Infections of SARS-CoV-2 were performed at a MOI of 1 for 24 hours, followed by drug treatment or 2% DMSO in triplicates. Uninfected cells were also treated with the same concentrations of drug or DMSO in triplicates. After 24 hours of drug treatment, total RNA from infected cells and uninfected cells were extracted using Qiagen RNeasy mini kit (Qiagen) following the manufacturer’s instructions. Then, we performed pair-end sequencing on a NovaSeq 6000 PE150 platform and generated 20 million reads per sample at Novogene Bioinformatics Institute (Novogene, Beijing, China).

### Bioinformatic Analyses

The raw reads quality was checked by the FastQC (0.11.7) and aligned to ChlSab1.1 (Chlorocebus sabaeus) reference genome by the <u>STAR</u> (2.5.0a) with default parameters. The count matrix was generated by the <u>featureCounts</u> (as a component of Subread package 2.0.1) program. Differentially expressed genes (DEGs) were calculated by <u>DESeq2</u> package (1.26.0) under R environment (3.6.1) and characterized for each sample (|L2FC| > 1, p-adjusted-value < 0.05). Gene set enrichment analysis (GSEA) was performed as previously described using normalized counts with orthology gene converting to human gene by <u>biomaRt</u> package (2.42.1)^50^. Bubble plots and heatmaps were generated using *ggplot* package and *heatmap()* function in R respectively.

### Statistical Analyses

For in vitro experiments, four-parameter dose response curve fitting was performed with constraints: Top = 1, IC_50_ > 0 using GraphPad Prism 8.

## Acknowledgments

We thank Dr. Martin Chan for helpful discussions during the early phase of this project. We thank Diana Grewe, Etienne Decroly, and Adrien Delpal for technical assistance. We also thank Prof. Vincent Lee and Dr. Ivanhoe Leung for the critical comments on this manuscript.

## Author contributions

W.L.N. & M.C.W.C. oversaw the project. H.S.L., H.-M.L., H.K., & W.L.N. wrote the manuscript with input from all authors. K.P.Y.H., K.-C.N. & J.C.W.H. contributed to the viral infection assay. H.S.L., S.K., H.M.L., J.H., B.S, C.E., J.-C.G., B.C., B.M. & W.S.A. contributed to the development of the RdRp assays. K.S.K., H.H.-Y.C. & K.B.W. contributed to the development of the M^pro^ assay. D.S., M.-H.L., K.P.W., I.D. & P.H.L. contributed to the development of the PL^pro^ assay. K.W. contributed to protein expression and purification. H.S.L., A.K.N.C., Y.W.C. & P.M.H.C. contributed to molecular modeling. Z.L. contributed to the mechanistic study. H.M.L., Z.Z., J.H., F.K.L.C., D.S.C.H., V.C.T.M. H.K., M.C.W.C. & W.L.N. contributed to the conceptual design of the project.

## Data and materials availability

The raw data and data analysis codes used in this project are available from the authors upon reasonable request.

---

**Supplementary Fig. 1.**
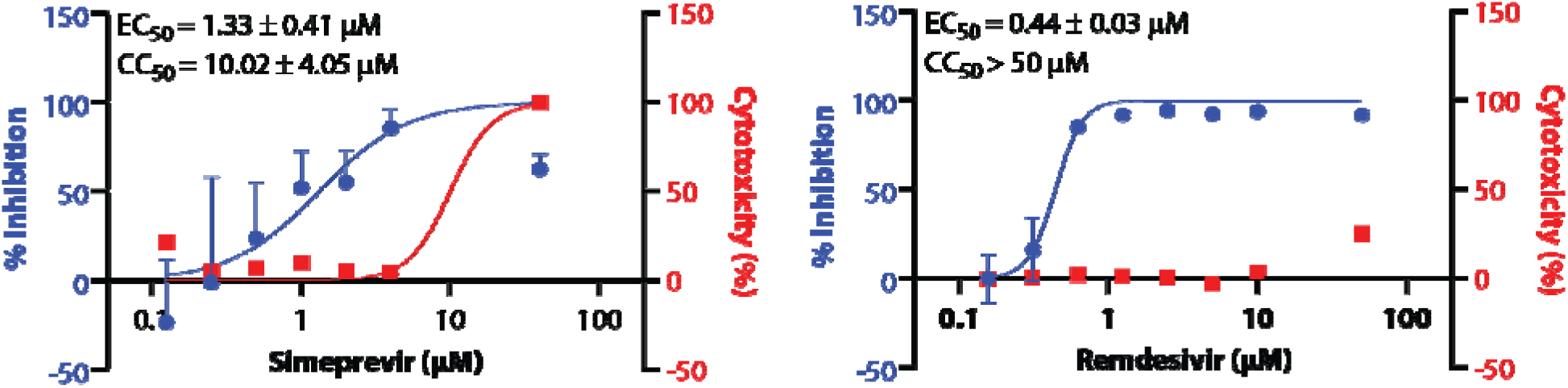
Dose-response curves in the suppression of SARS-CoV-2 replication in A549-ACE2 cells and cytotoxicity for simeprevir **(Left)** and remdesivir **(Right)** are shown. Data points in all plots represent mean ± S.E.M‥ For all data points, *n* = 3 replicates.

**Supplementary Fig. 2.**
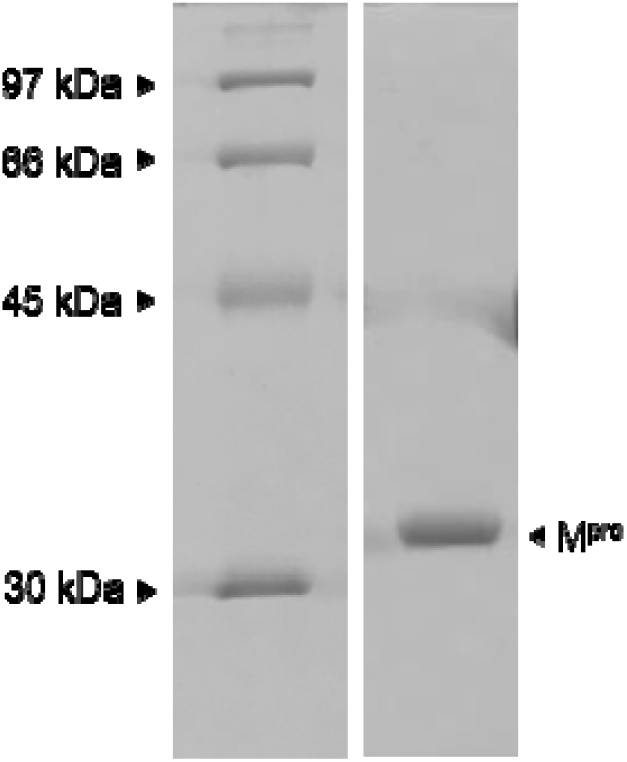
SDS-PAGE analysis of recombinant SARS-CoV-2 M^pro^.

**Supplementary Fig. 3.**
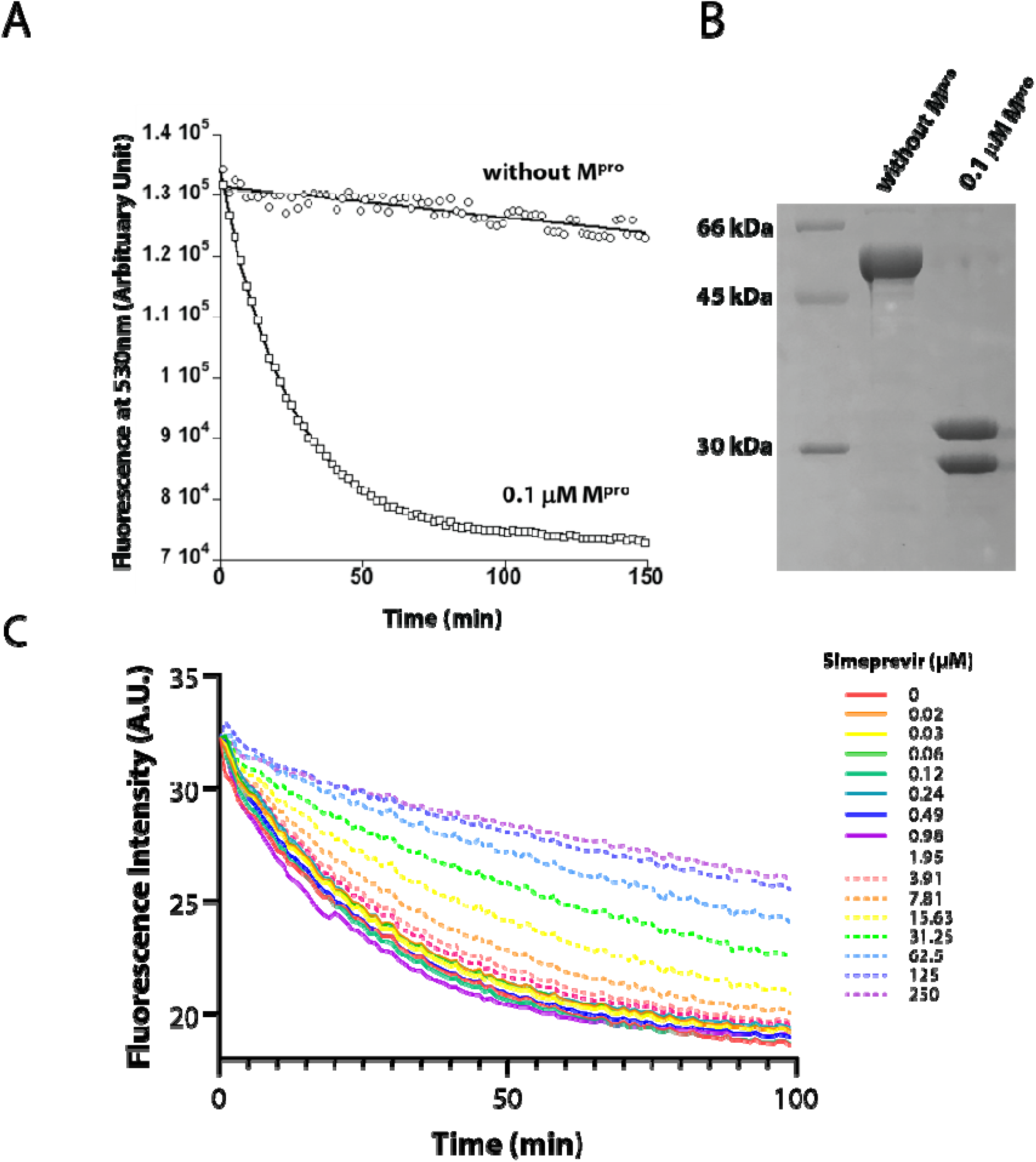
**(A)** The addition of SARS-CoV-2 M^pro^ led to the cleavage of the substrate, causing detectable decline in FRET signal with 430-nm excitation and 530-nm emission. **(B)** Confirmation of substrate cleavage by M^pro^ using SDS-PAGE. **(C)** The addition of simeprevir at varying concentrations attenuated the rate of FRET substrate cleavage by M^pro^.

**Supplementary Fig. 4.**
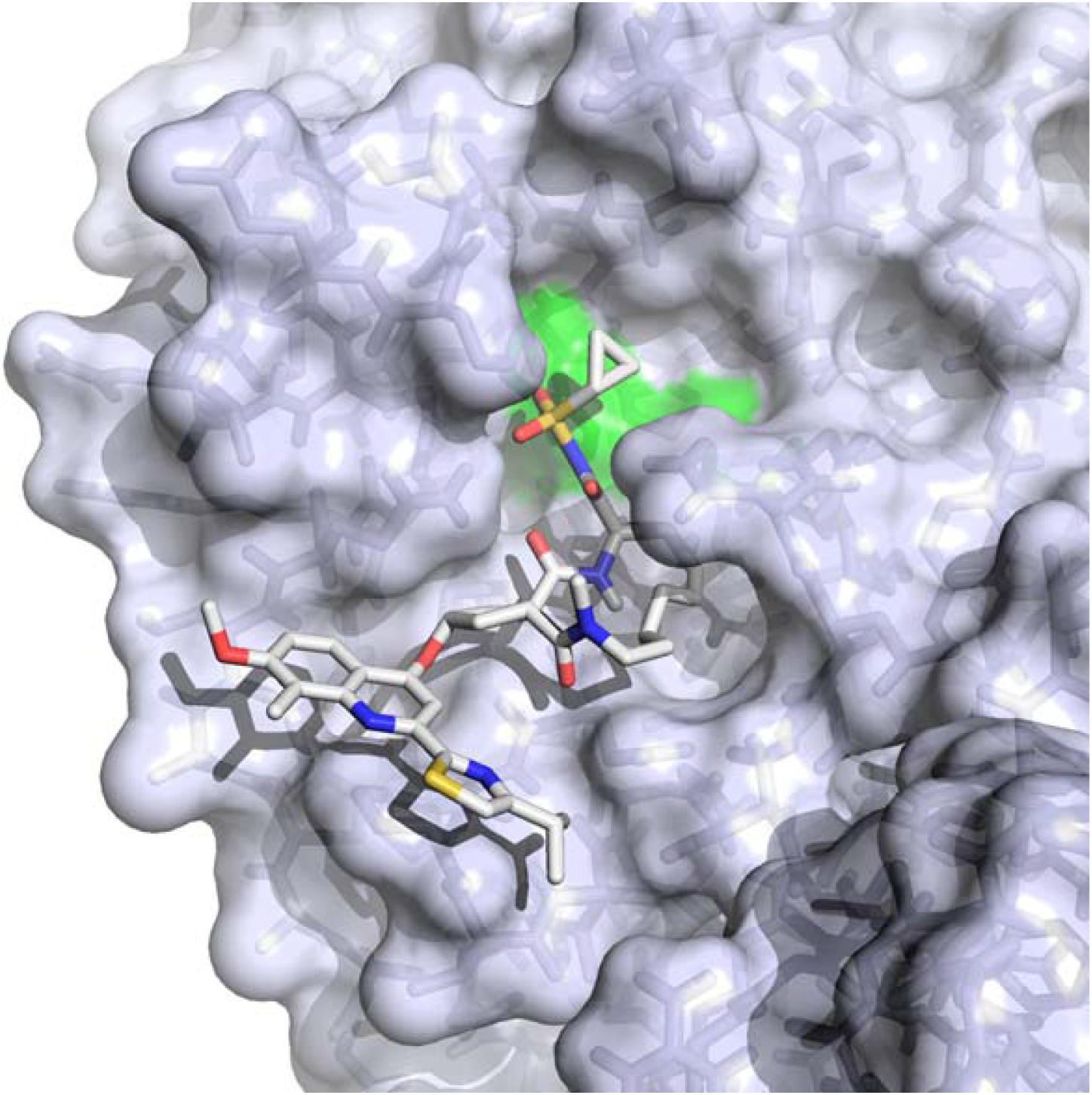
Docking of simeprevir on SARS-CoV-2 M^pro^ (performed with AutoDock Vina version 1.1.2). The M^pro^ structure was based on an apo protein crystal structure (PDB ID: 6YB7); the A:B dimer was generated by crystallographic symmetry. Docking was run with the substrate-binding residues set to be flexible; and a 30 × 30 × 30 Å^3^ search box centered near the side-chain N∊2 atom of His163. The top docking mode shown here scored −9.9 kcal mol^−1^. The protein is shown as a semi-transparent molecular surface encasing its stick model with the catalytic residues His41 and Cys145 in green. The ligand is shown as a stick model with oxygen atoms in red; nitrogen atoms in blue, sulfur atoms in yellow and carbon atoms in white.

**Supplementary Fig. 5.**
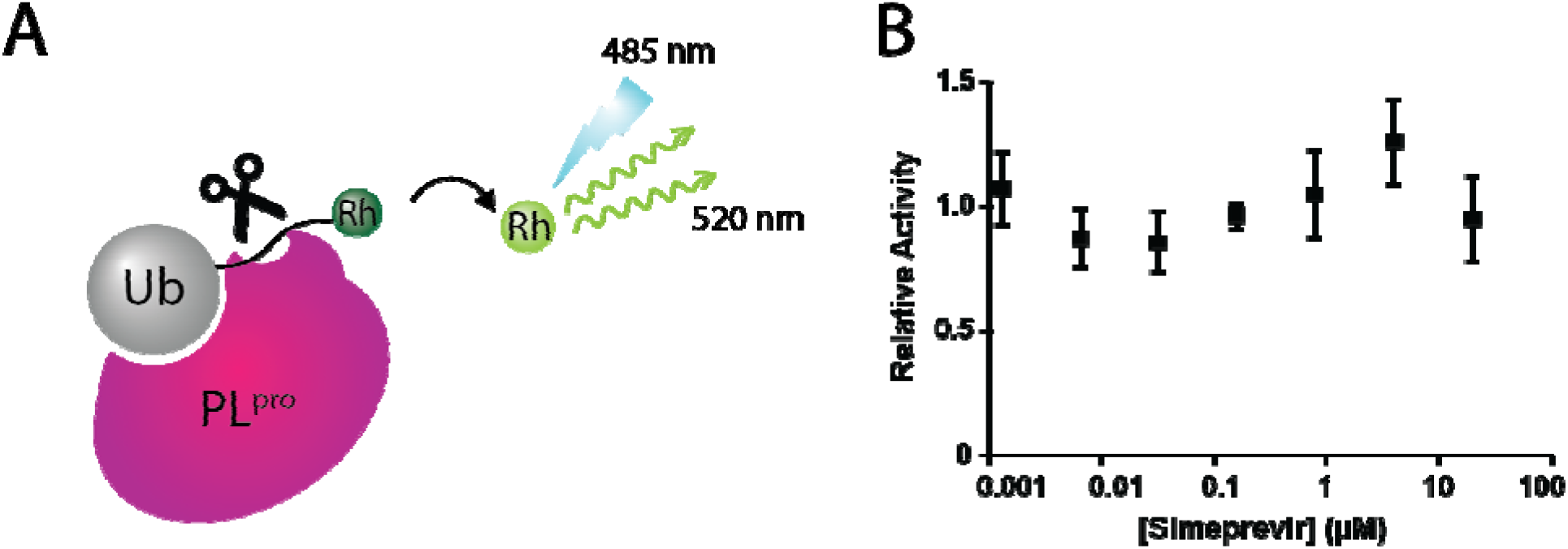
Simeprevir did not inhibit PL^pro^ activity in a repeated PL^pro^ cleavage assay using a different substrate ubiquitin-rhodamine.

**Supplementary Fig. 6.**
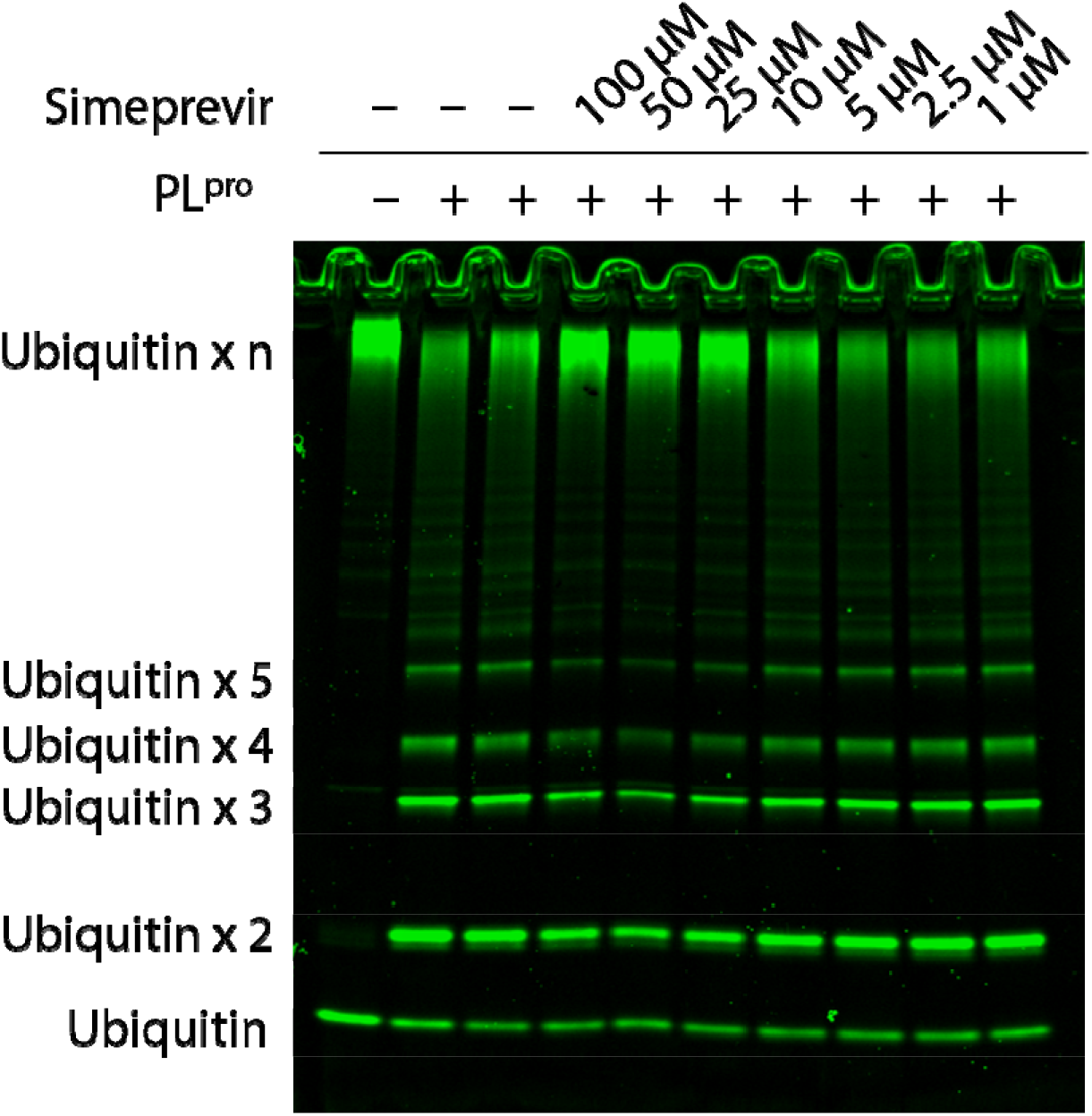
Inhibitor deubiquitination assay of PL^pro^ shows PL^pro^ (SARS-CoV-2) actively cleaves poly-ubiquitinated substrate into di-ubiquitin or longer ubiquitin chain. While simeprevir was added in reaction solution, the activity of PL^pro^ was not profoundly inhibited compared to control experiments (no inhibitor and no inhibitor with 2% DMSO).

**Supplementary Fig. 7.**
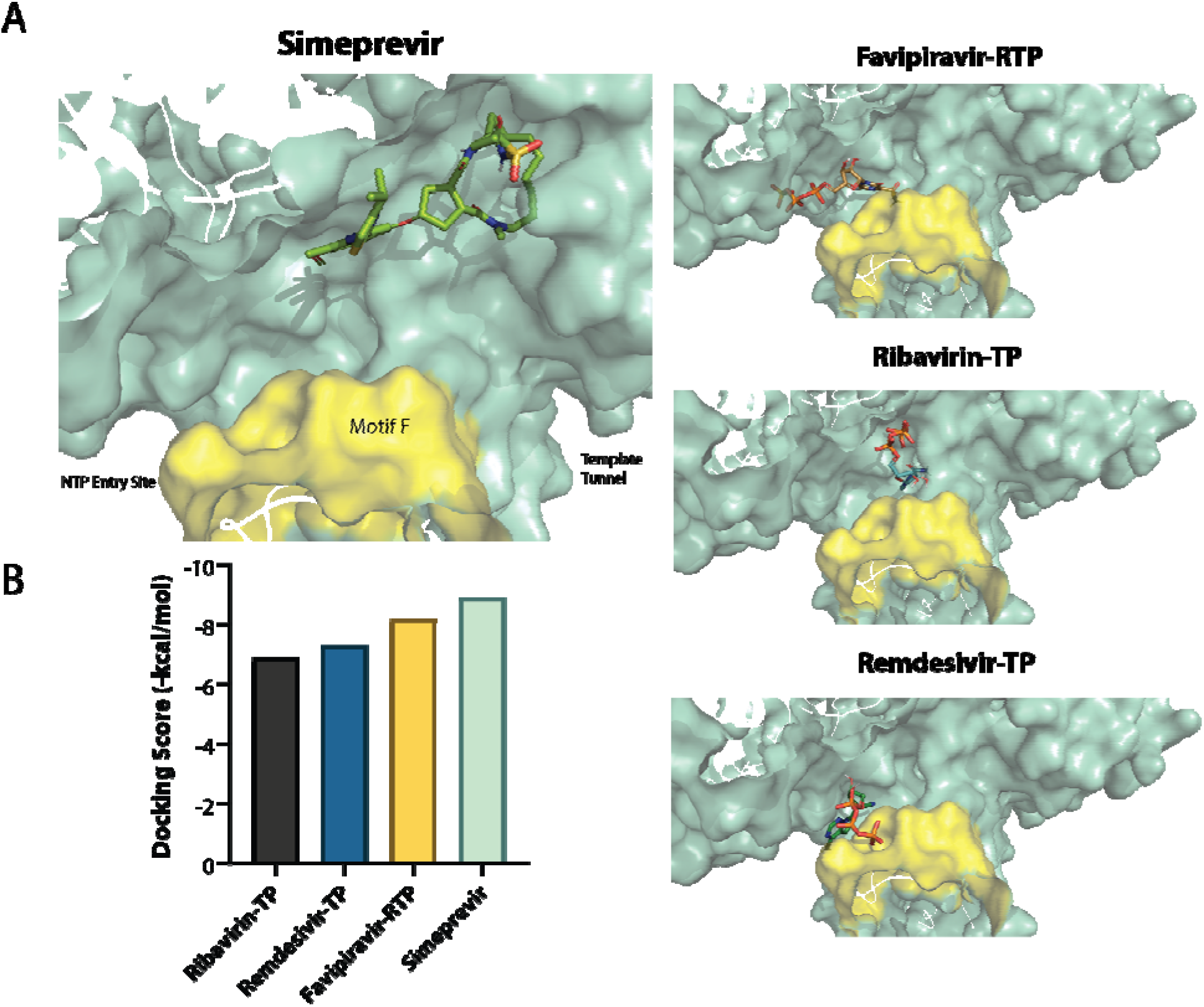
**(A)** Binding mode of simeprevir and other inhibitors against SARS-CoV-2 nsp12 (PDB ID: 6M71). Motif F is highlighted in yellow. **(B)** Docking scores of drug candidates against nsp12.

**Supplementary Fig. 8.**
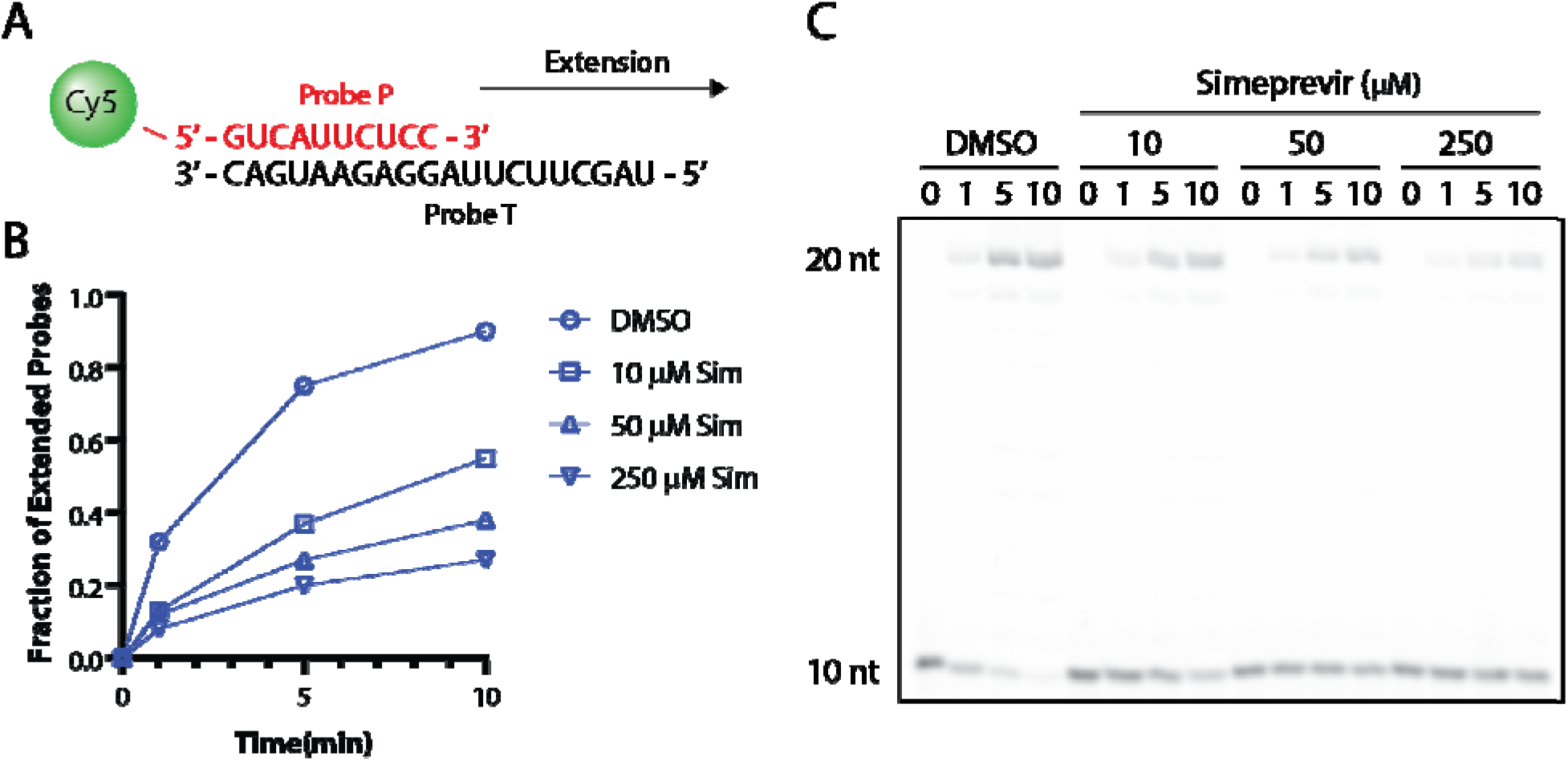
**(A)** Gel-based assay scheme for SARS-CoV-RdRp. Two partially complementary RNA probes are extended by the enzyme, and the extent of extension is visualized by detecting the Cy5 labeling of the shorter probe P. **(B)** Time-dependent elongation of probe P. Fraction of extension is determined by densitometry of extended product versus total RNA. **(C)** RNA-PAGE imaged with Cy5 mode.

**Supplementary Fig. 9.**
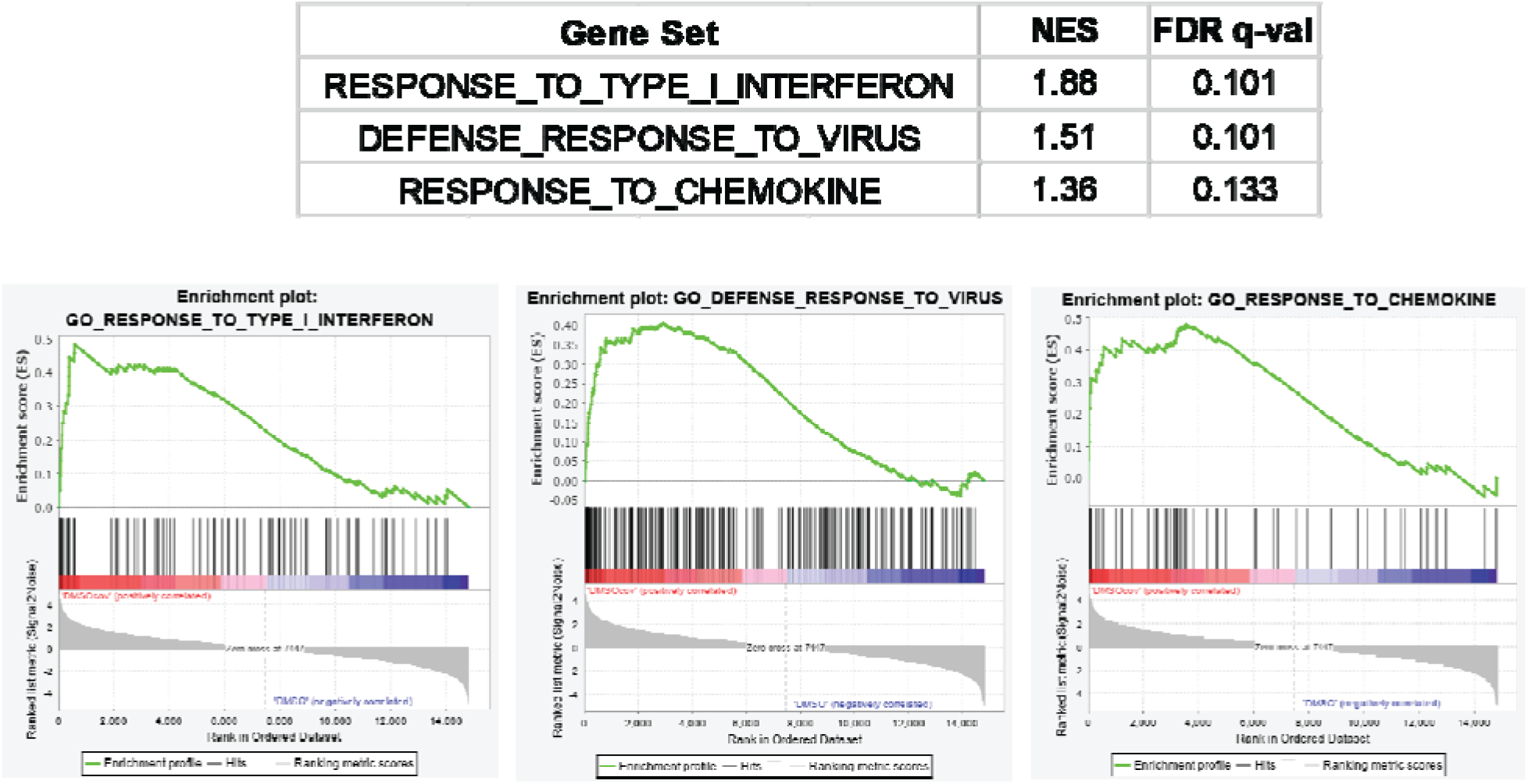
Gene set enrichment analysis of SARS-CoV-2 infection. Relevant statistics and enrichment plots for three gene sets in response to virus infection are shown.

**Supplementary Table 1.** FDA-approved repurposable drug candidates tested in this study.

| **Drug candidate** | **Approved indication(s)** | **Common/notable adverse effects** | **Drug formulation(s) available** | **Contraindications (except known hypersensitivity to the drug)** |
| --- | --- | --- | --- | --- |
| Simeprevir | Chronic HCV infection | Photosensitivity, rash, fatigue, myalgia, dyspnea | Oral | None |
| Daclatasvir | Chronic HCV infection | Fatigue, nausea, anemia, diarrhea | Oral | None |
| Saquinavir | HIV infection | Nausea, diarrhea, QT prolongation | Oral | Prolonged QT interval, Patients at risk/having complete AV block |
| Bromocriptine | Hyperprolactinemia, pituitary prolactinoma | Constipation, dizziness, nausea, fatigue, orthostatic hypotension, vasospasm, abdominal pain | Oral | Uncontrolled hypertension, psychosis, syncopal migraine |
| Asunaprevir | HIV infection | Fatigue, rash, nausea, neutropenia, anemia, deranged liver function tests | Oral | Moderate/severe hepatic impairment (Child B or C) |
| Bictegravir | HIV infection | Increased serum creatine kinase, deranged liver function tests, neutropenia | Oral | None |
| Entecavir | HBV infection | Deranged liver function tests | Oral | Moderate/severe hepatic impairment (Child B or C) |
| Zidovudine | HIV infection | Headache, malaise, rash, nausea, neutropenia, anemia | Oral | None |
| Sofosbuvir | Chronic HCV infection | Fatigue, headache, insomnia, nausea, diarrhea, anemia, myalgia, rash | Oral | None |
| Atovaquone | Protozoal infection or prophylaxis (Pneumocystis jirovecii, Plasmodium spp. etc) | Headache, insomnia, rash, diarrhea, myalgia, drug fever, hyponatremia, neutropenia | Oral | None |
| Remdesivir | COVID-19 | Deranged liver function tests | Intravenous | Serum alanine transaminase $\geq 5$x the upper-limit of normal |

**Supplementary Table 2.**
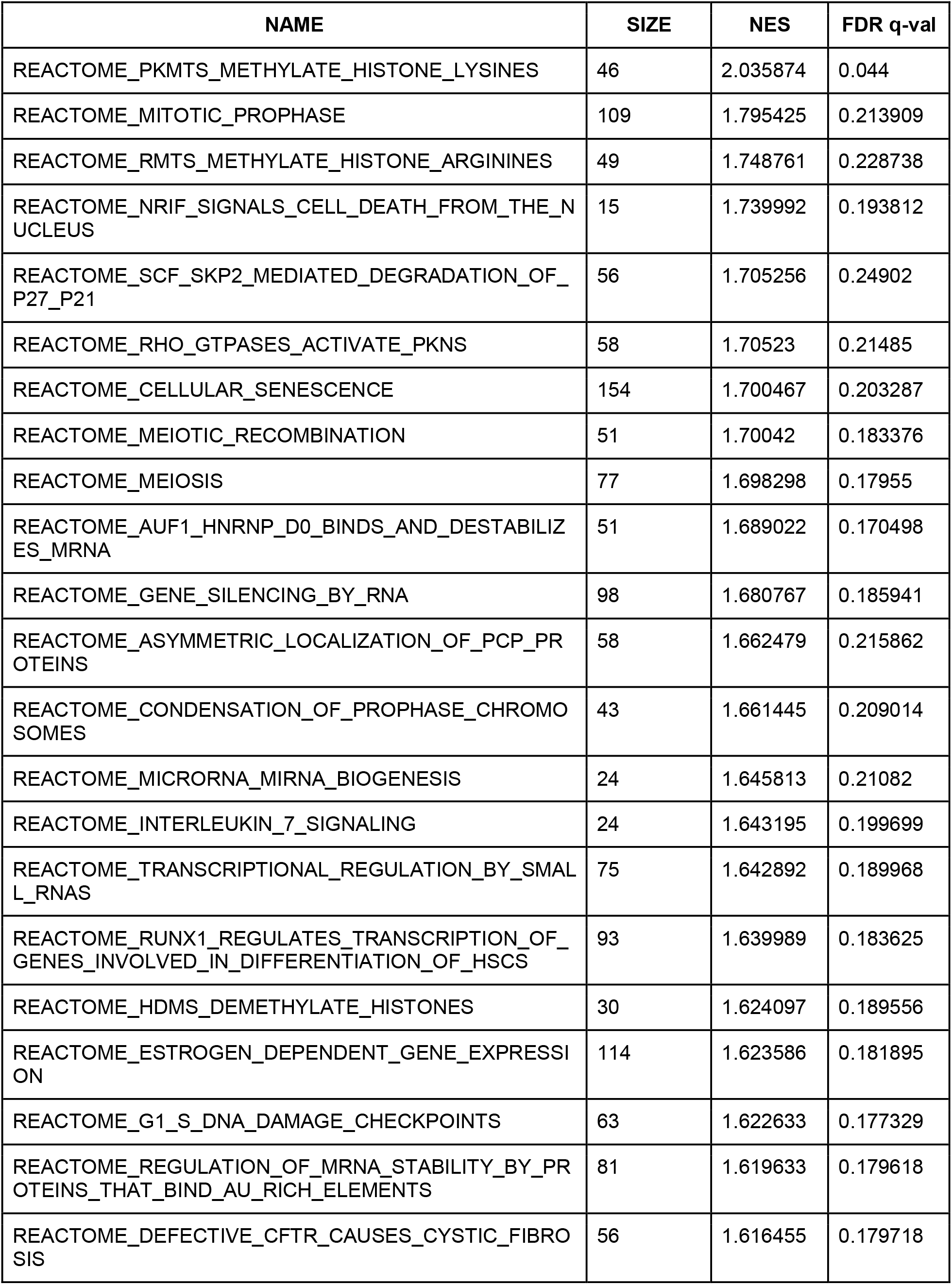
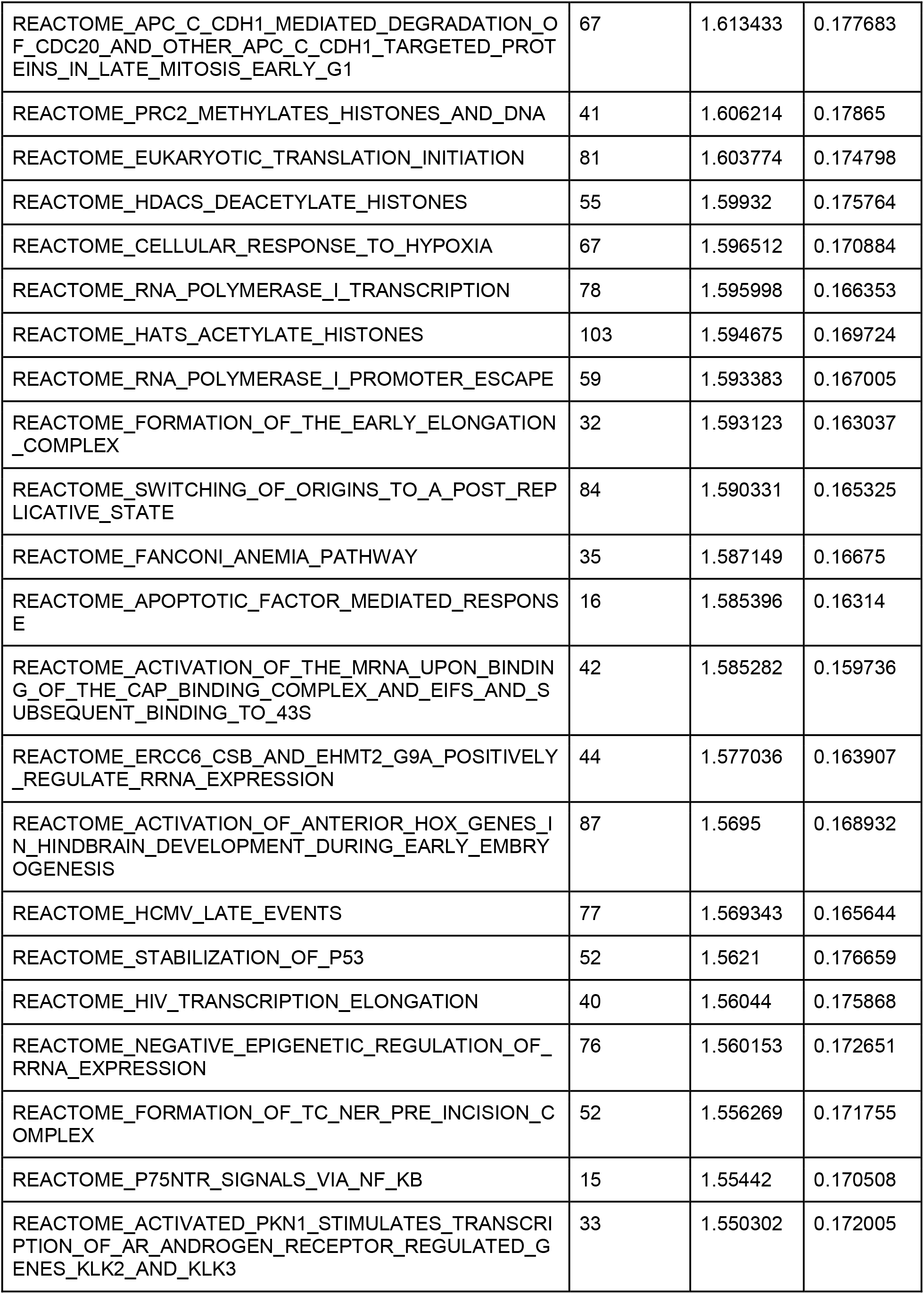
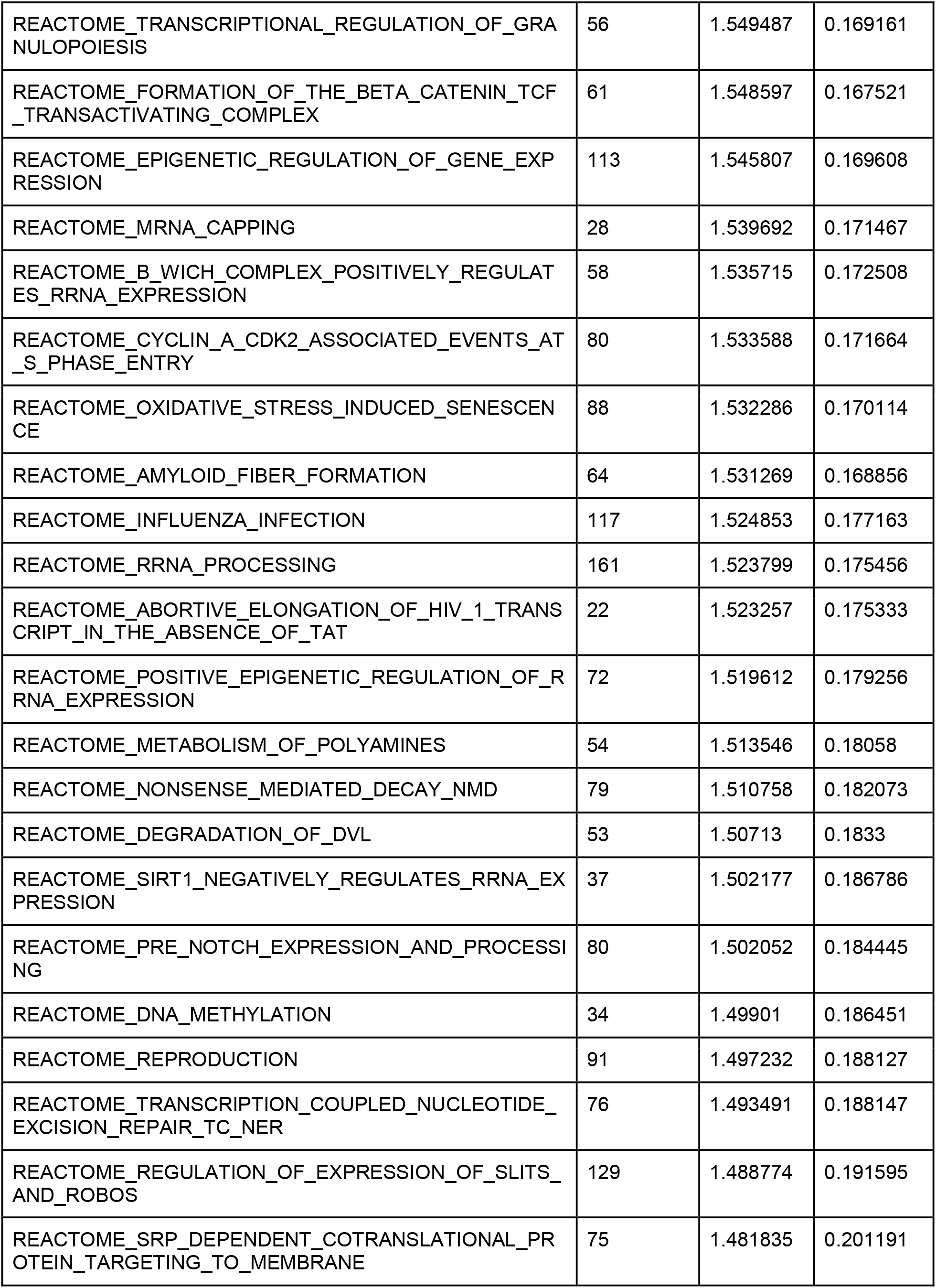
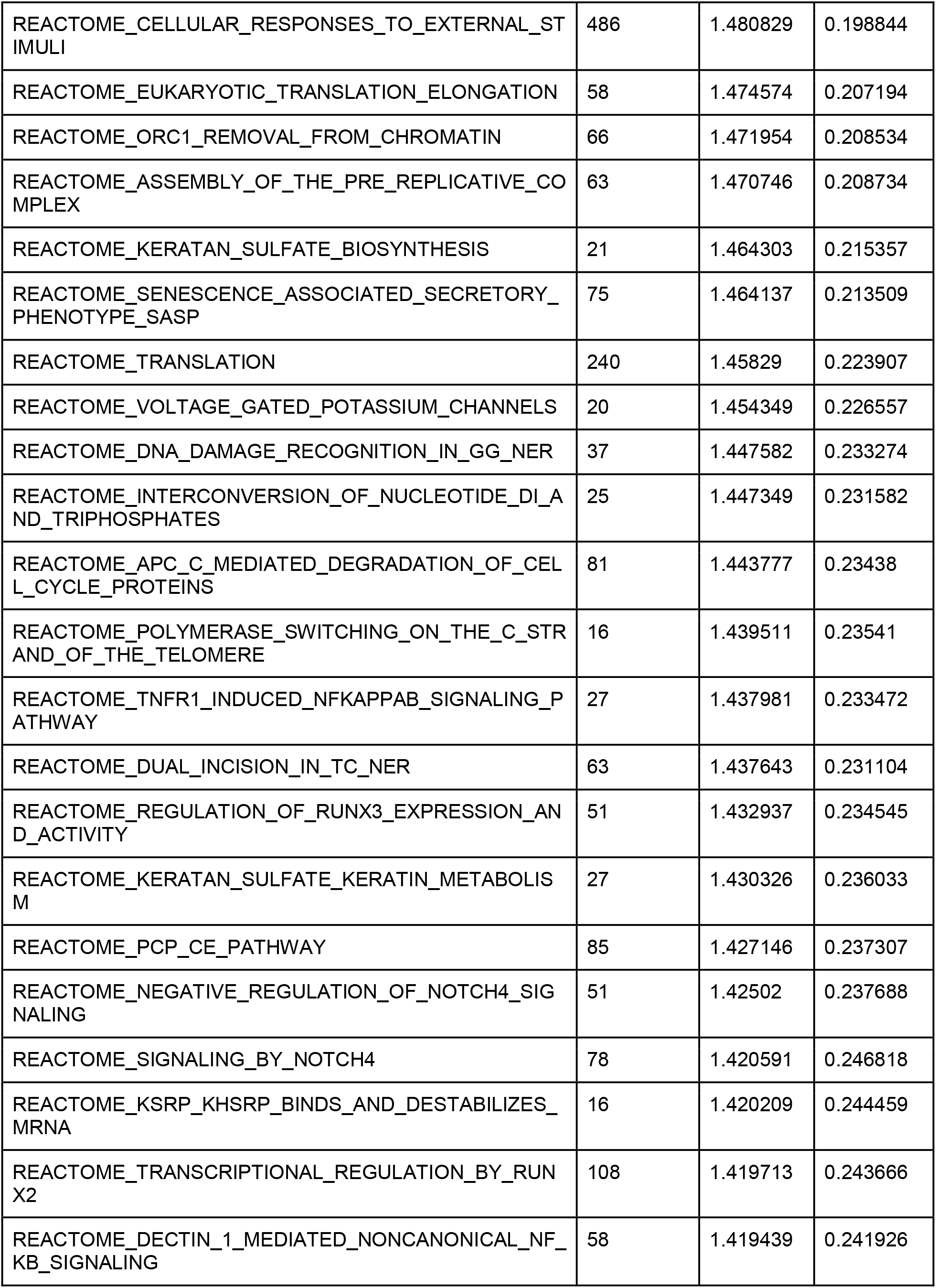
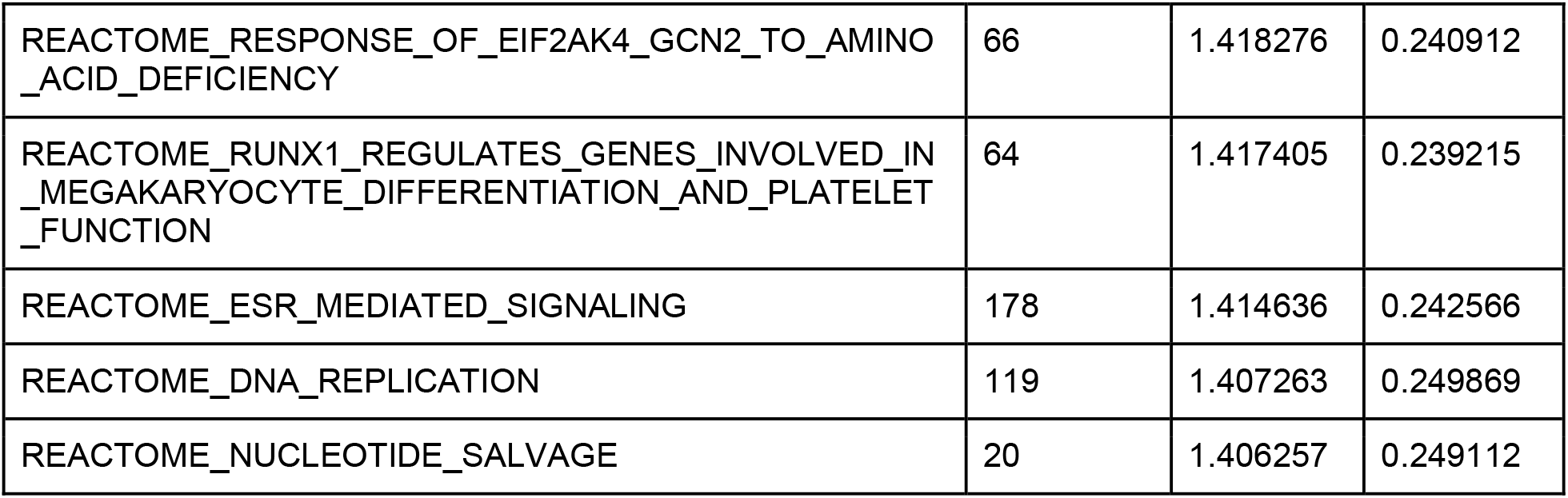
Gene sets enriched in Sim 1.1 vs. DMSO in infected samples according to Gene set enrichment analysis (GSEA).

## Reference

(1) Huang, C.; Wang, Y.; Li, X.; Ren, L.; Zhao, J.; Hu, Y.; Zhang, L.; Fan, G.; Xu, J.; Gu, X.; et al. Clinical Features of Patients Infected with 2019 Novel Coronavirus in Wuhan, China. Lancet 2020, 395 (10223), 497–506.

(2) Chen, N.; Zhou, M.; Dong, X.; Qu, J.; Gong, F.; Han, Y.; Qiu, Y.; Wang, J.; Liu, Y.; Wei, Y.; et al. Epidemiological and Clinical Characteristics of 99 Cases of 2019 Novel Coronavirus Pneumonia in Wuhan, China: A Descriptive Study. Lancet 2020, 395 (10223), 507–513.

(3) Zhou, F.; Yu, T.; Du, R.; Fan, G.; Liu, Y.; Liu, Z.; Xiang, J.; Wang, Y.; Song, B.; Gu, X.; et al. Clinical Course and Risk Factors for Mortality of Adult Inpatients with COVID-19 in Wuhan, China: A Retrospective Cohort Study. Lancet 2020, 395 (10229), 1054–1062.

(4) Guan, W.-J.; Ni, Z.-Y.; Hu, Y.; Liang, W.-H.; Ou, C.-Q.; He, J.-X.; Liu, L.; Shan, H.; Lei, C.-L.; Hui, D. S. C.; et al. Clinical Characteristics of Coronavirus Disease 2019 in China. N. Engl. J. Med. 2020.

(5) Li, R.; Pei, S.; Chen, B.; Song, Y.; Zhang, T.; Yang, W.; Shaman, J. Substantial Undocumented Infection Facilitates the Rapid Dissemination of Novel Coronavirus (SARS-CoV-2). Science 2020, 368 (6490), 489–493.

(6) Sanders, J. M.; Monogue, M. L.; Jodlowski, T. Z.; Cutrell, J. B. Pharmacologic Treatments for Coronavirus Disease 2019 (COVID-19): A Review. JAMA 2020.

(7) Pastick, K. A.; Okafor, E. C.; Wang, F.; Lofgren, S. M.; Skipper, C. P.; Nicol, M. R.; Pullen, M. F.; Rajasingham, R.; McDonald, E. G.; Lee, T. C.; et al. Review: Hydroxychloroquine and Chloroquine for Treatment of SARS-CoV-2 (COVID-19). Open Forum Infect. Dis. 2020, 7 (4), ofaa130.

(8) Zumla, A.; Chan, J. F. W.; Azhar, E. I.; Hui, D. S. C.; Yuen, K.-Y. Coronaviruses - Drug Discovery and Therapeutic Options. Nat. Rev. Drug Discov. 2016, 15 (5), 327–347.

(9) Sheahan, T. P.; Sims, A. C.; Zhou, S.; Graham, R. L.; Pruijssers, A. J.; Agostini, M. L.; Leist, S. R.; Schäfer, A.; Dinnon, K. H.; Stevens, L. J.; et al. An Orally Bioavailable Broad-Spectrum Antiviral Inhibits SARS-CoV-2 in Human Airway Epithelial Cell Cultures and Multiple Coronaviruses in Mice. Sci. Transl. Med. 2020, 12 (541).

(10) Gordon, C. J.; Tchesnokov, E. P.; Woolner, E.; Perry, J. K.; Feng, J. Y.; Porter, D. P.; Götte, M. Remdesivir Is a Direct-Acting Antiviral That Inhibits RNA-Dependent RNA Polymerase from Severe Acute Respiratory Syndrome Coronavirus 2 with High Potency. J. Biol. Chem. 2020, 295 (20), 6785–6797.

(11) Yin, W.; Mao, C.; Luan, X.; Shen, D.-D.; Shen, Q.; Su, H.; Wang, X.; Zhou, F.; Zhao, W.; Gao, M.; et al. Structural Basis for Inhibition of the RNA-Dependent RNA Polymerase from SARS-CoV-2 by Remdesivir. Science 2020, 368 (6498), 1499–1504.

(12) Rosenberg, E. S.; Dufort, E. M.; Udo, T.; Wilberschied, L. A.; Kumar, J.; Tesoriero, J.; Weinberg, P.; Kirkwood, J.; Muse, A.; DeHovitz, J.; et al. Association of Treatment With Hydroxychloroquine or Azithromycin With In-Hospital Mortality in Patients With COVID-19 in New York State. JAMA 2020.

(13) Geleris, J.; Sun, Y.; Platt, J.; Zucker, J.; Baldwin, M.; Hripcsak, G.; Labella, A.; Manson, D. K.; Kubin, C.; Barr, R. G.; et al. Observational Study of Hydroxychloroquine in Hospitalized Patients with Covid-19. N. Engl. J. Med. 2020, 382 (25), 2411–2418.

(14) Cao, B.; Wang, Y.; Wen, D.; Liu, W.; Wang, J.; Fan, G.; Ruan, L.; Song, B.; Cai, Y.; Wei, M.; et al. A Trial of Lopinavir-Ritonavir in Adults Hospitalized with Severe Covid-19. N. Engl. J. Med. 2020, 382 (19), 1787–1799.

(15) Gbinigie, K.; Frie, K. Should Chloroquine and Hydroxychloroquine Be Used to Treat COVID-19? A Rapid Review. Br J Gen Pract Open 2020, 4 (2).

(16) Chorin, E.; Wadhwani, L.; Magnani, S.; Dai, M.; Shulman, E.; Nadeau-Routhier, C.; Knotts, R.; Bar-Cohen, R.; Kogan, E.; Barbhaiya, C.; et al. QT Interval Prolongation and Torsade de Pointes in Patients with COVID-19 Treated with Hydroxychloroquine/Azithromycin. Heart Rhythm 2020.

(17) Wang, Y.; Zhang, D.; Du, G.; Du, R.; Zhao, J.; Jin, Y.; Fu, S.; Gao, L.; Cheng, Z.; Lu, Q.; et al. Remdesivir in Adults with Severe COVID-19: A Randomised, Double-Blind, Placebo-Controlled, Multicentre Trial. Lancet 2020, 395 (10236), 1569–1578.

(18) Ko, M.; Jeon, S.; Ryu, W.-S.; Kim, S. Comparative Analysis of Antiviral Efficacy of FDA-Approved Drugs against SARS-CoV-2 in Human Lung Cells: Nafamostat Is the Most Potent Antiviral Drug Candidate. BioRxiv 2020.

(19) Weston, S.; Haupt, R.; Logue, J.; Matthews, K.; Frieman, M. FDA Approved Drugs with Broad Anti-Coronaviral Activity Inhibit SARS-CoV-2 *in Vitro*. BioRxiv 2020.

(20) Wang, M.; Cao, R.; Zhang, L.; Yang, X.; Liu, J.; Xu, M.; Shi, Z.; Hu, Z.; Zhong, W.; Xiao, G. Remdesivir and Chloroquine Effectively Inhibit the Recently Emerged Novel Coronavirus (2019-NCoV) in Vitro. Cell Res. 2020, 30 (3), 269–271.

(21) Choy, K.-T.; Wong, A. Y.-L.; Kaewpreedee, P.; Sia, S. F.; Chen, D.; Hui, K. P. Y.; Chu, D. K. W.; Chan, M. C. W.; Cheung, P. P.-H.; Huang, X.; et al. Remdesivir, Lopinavir, Emetine, and Homoharringtonine Inhibit SARS-CoV-2 Replication in Vitro. Antiviral Res. 2020, 178, 104786.

(22) Riva, L.; Yuan, S.; Yin, X.; Martin-Sancho, L.; Matsunaga, N.; Pache, L.; Burgstaller-Muehlbacher, S.; De Jesus, P. D.; Teriete, P.; Hull, M. V.; et al. Discovery of SARS-CoV-2 Antiviral Drugs through Large-Scale Compound Repurposing. Nature 2020.

(23) Zhang, L.; Lin, D.; Sun, X.; Curth, U.; Drosten, C.; Sauerhering, L.; Becker, S.; Rox, K.; Hilgenfeld, R. Crystal Structure of SARS-CoV-2 Main Protease Provides a Basis for Design of Improved α-Ketoamide Inhibitors. Science 2020, 368 (6489), 409–412.

(24) Jin, Z.; Du, X.; Xu, Y.; Deng, Y.; Liu, M.; Zhao, Y.; Zhang, B.; Li, X.; Zhang, L.; Peng, C.; et al. Structure of Mpro from SARS-CoV-2 and Discovery of Its Inhibitors. Nature 2020.

(25) Shin, D.; Mukherjee, R.; Grewe, D.; Bojkova, D.; Baek, K.; Bhattacharya, A.; Schulz, L.; Widera, M.; Mehdipour, A. R.; Tascher, G.; et al. Inhibition of Papain-like Protease PLpro Blocks SARS-CoV-2 Spread and Promotes Anti-Viral Immunity. 2020.

(26) Gao, Y.; Yan, L.; Huang, Y.; Liu, F.; Zhao, Y.; Cao, L.; Wang, T.; Sun, Q.; Ming, Z.; Zhang, L.; et al. Structure of the RNA-Dependent RNA Polymerase from COVID-19 Virus. Science 2020, 368 (6492), 779–782.

(27) Jin, Z.; Zhao, Y.; Sun, Y.; Zhang, B.; Wang, H.; Wu, Y.; Zhu, Y.; Zhu, C.; Hu, T.; Du, X.; et al. Structural Basis for the Inhibition of SARS-CoV-2 Main Protease by Antineoplastic Drug Carmofur. Nat. Struct. Mol. Biol. 2020.

(28) Rosenquist, Å.; Samuelsson, B.; Johansson, P.-O.; Cummings, M. D.; Lenz, O.; Raboisson, P.; Simmen, K.; Vendeville, S.; de Kock, H.; Nilsson, M.; et al. Discovery and Development of Simeprevir (TMC435), a HCV NS3/4A Protease Inhibitor. J. Med. Chem. 2014, 57 (5), 1673–1693.

(29) Mossel, E. C.; Huang, C.; Narayanan, K.; Makino, S.; Tesh, R. B.; Peters, C. J. Exogenous ACE2 Expression Allows Refractory Cell Lines to Support Severe Acute Respiratory Syndrome Coronavirus Replication. J. Virol. 2005, 79 (6), 3846–3850.

(30) Lin, T.-I.; Lenz, O.; Fanning, G.; Verbinnen, T.; Delouvroy, F.; Scholliers, A.; Vermeiren, K.; Rosenquist, A.; Edlund, M.; Samuelsson, B.; et al. In Vitro Activity and Preclinical Profile of TMC435350, a Potent Hepatitis C Virus Protease Inhibitor. Antimicrob. Agents Chemother. 2009, 53 (4), 1377–1385.

(31) Chuck, C.-P.; Chong, L.-T.; Chen, C.; Chow, H.-F.; Wan, D. C.-C.; Wong, K.-B. Profiling of Substrate Specificity of SARS-CoV 3CL. PLoS ONE 2010, 5 (10), e13197.

(32) Ma, C.; Hurst, B.; Hu, Y.; Szeto, T.; Tarbet, B.; Wang, J. Boceprevir, GC-376, and Calpain Inhibitors II, XII Inhibit SARS-CoV-2 Viral Replication by Targeting the Viral Main Protease. BioRxiv 2020.

(33) Calligari, P.; Bobone, S.; Ricci, G.; Bocedi, A. Molecular Investigation of SARS-CoV-2 Proteins and Their Interactions with Antiviral Drugs. Viruses 2020, 12 (4).

(34) Subissi, L.; Posthuma, C. C.; Collet, A.; Zevenhoven-Dobbe, J. C.; Gorbalenya, A. E.; Decroly, E.; Snijder, E. J.; Canard, B.; Imbert, I. One Severe Acute Respiratory Syndrome Coronavirus Protein Complex Integrates Processive RNA Polymerase and Exonuclease Activities. Proc Natl Acad Sci USA 2014, 111 (37), E3900–9.

(35) Blanco-Melo, D.; Nilsson-Payant, B. E.; Liu, W.-C.; Uhl, S.; Hoagland, D.; Møller, R.; Jordan, T. X.; Oishi, K.; Panis, M.; Sachs, D.; et al. Imbalanced Host Response to SARS-CoV-2 Drives Development of COVID-19. Cell 2020, 181 (5), 1036–1045.e9.

(36) Park, A.; Iwasaki, A. Type I and Type III Interferons - Induction, Signaling, Evasion, and Application to Combat COVID-19. Cell Host Microbe 2020, 27 (6), 870–878.

(37) Janssen Pharmaceutica. Clinical Pharmacology and Biopharmaceutics Reivew(s) (Application Number 205123Orig1s000), Center for Drug Evaluation and Research.

(38) Snoeys, J.; Beumont, M.; Monshouwer, M.; Ouwerkerk-Mahadevan, S. Mechanistic Understanding of the Nonlinear Pharmacokinetics and Intersubject Variability of Simeprevir: A PBPK-Guided Drug Development Approach. Clin. Pharmacol. Ther. 2016, 99 (2), 224–234.

(39) Ma, C.; Sacco, M. D.; Hurst, B.; Townsend, J. A.; Hu, Y.; Szeto, T.; Zhang, X.; Tarbet, B.; Marty, M. T.; Chen, Y.; et al. Boceprevir, GC-376, and Calpain Inhibitors II, XII Inhibit SARS-CoV-2 Viral Replication by Targeting the Viral Main Protease. Cell Res. 2020, 30 (8), 678–692.

(40) Wei, J.; Alfajaro, M. M.; Hanna, R. E.; DeWeirdt, P. C.; Strine, M. S.; Lu-Culligan, W. J.; Zhang, S. M.; Graziano, V. R.; Schmitz, C. O.; Chen, J. S.; et al. Genome-Wide CRISPR Screen Reveals Host Genes That Regulate SARS-CoV-2 Infection. BioRxiv 2020.

(41) Marcos-Villar, L.; Díaz-Colunga, J.; Sandoval, J.; Zamarreño, N.; Landeras-Bueno, S.; Esteller, M.; Falcón, A.; Nieto, A. Epigenetic Control of Influenza Virus: Role of H3K79 Methylation in Interferon-Induced Antiviral Response. Sci. Rep. 2018, 8 (1), 1230.

(42) Menachery, V. D.; Schäfer, A.; Burnum-Johnson, K. E.; Mitchell, H. D.; Eisfeld, A. J.; Walters, K. B.; Nicora, C. D.; Purvine, S. O.; Casey, C. P.; Monroe, M. E.; et al. MERS-CoV and H5N1 Influenza Virus Antagonize Antigen Presentation by Altering the Epigenetic Landscape. Proc Natl Acad Sci USA 2018, 115 (5), E1012–E1021.

(43) Hui, K. P. Y.; Cheung, M.-C.; Perera, R. A. P. M.; Ng, K.-C.; Bui, C. H. T.; Ho, J. C. W.; Ng, M. M. T.; Kuok, D. I. T.; Shih, K. C.; Tsao, S.-W.; et al. Tropism, Replication Competence, and Innate Immune Responses of the Coronavirus SARS-CoV-2 in Human Respiratory Tract and Conjunctiva: An Analysis in Ex-Vivo and in-Vitro Cultures. Lancet Respir. Med. 2020.

(44) Sterling, T.; Irwin, J. J. ZINC 15--Ligand Discovery for Everyone. J. Chem. Inf. Model. 2015, 55 (11), 2324–2337.

(45) Trott, O.; Olson, A. J. AutoDock Vina: Improving the Speed and Accuracy of Docking with a New Scoring Function, Efficient Optimization, and Multithreading. J. Comput. Chem. 2010, 31 (2), 455–461.

(46) Chuck, C.-P.; Chen, C.; Ke, Z.; Wan, D. C.-C.; Chow, H.-F.; Wong, K.-B. Design, Synthesis and Crystallographic Analysis of Nitrile-Based Broad-Spectrum Peptidomimetic Inhibitors for Coronavirus 3C-like Proteases. Eur. J. Med. Chem. 2013, 59, 1–6.

(47) Shannon, A.; Selisko, B.; Le, N.; Huchting, J.; Touret, F.; Piorkowski, G.; Fattorini, V.; Ferron, F.; Decroly, E.; Meier, C.; et al. Favipiravir Strikes the SARS-CoV-2 at Its Achilles Heel, the RNA Polymerase. BioRxiv 2020.

(48) Eydoux, C.; Fattorini, V.; Shannon, A.; Le, T.-T.-N.; Didier, B.; Canard, B.; Guillemot, J.-C. A Fluorescence-Based High Throughput-Screening Assay for the SARS-CoV RNA Synthesis Complex. BioRxiv 2020.

(49) Zhang, W.; Wu, K.-P.; Sartori, M. A.; Kamadurai, H. B.; Ordureau, A.; Jiang, C.; Mercredi, P. Y.; Murchie, R.; Hu, J.; Persaud, A.; et al. System-Wide Modulation of HECT E3 Ligases with Selective Ubiquitin Variant Probes. Mol. Cell 2016, 62 (1), 121–136.

(50) Subramanian, A.; Tamayo, P.; Mootha, V. K.; Mukherjee, S.; Ebert, B. L.; Gillette, M. A.; Paulovich, A.; Pomeroy, S. L.; Golub, T. R.; Lander, E. S.; et al. Gene Set Enrichment Analysis: A Knowledge-Based Approach for Interpreting Genome-Wide Expression Profiles. Proc Natl Acad Sci USA 2005, 102 (43), 15545–15550.

